# The complex swarming dynamics of malaria mosquitoes emerge from simple minimally-interactive behavioral rules

**DOI:** 10.1101/2024.08.31.610631

**Authors:** Antoine Cribellier, Bèwadéyir Serge Poda, Roch K. Dabiré, Abdoulaye Diabaté, Olivier Roux, Florian T. Muijres

**Affiliations:** Experimental Zoology Group, Wageningen University, Wageningen, The Netherlands; Institut de Recherche en Sciences de la Santé (IRSS), Bobo-Dioulasso, Burkina Faso; MIVEGEC, University of Montpellier, IRD, CNRS, Montpellier, France

## Abstract

Swarming is a widespread collective behavior in animals, often thought to emerge from complex interactions among individuals. Here, we show that mating swarms of malaria mosquitoes can emerge from simple behavioral rules with minimal interaction between individuals. We analyzed two published experimental datasets with three-dimensional flight tracks of *Anopheles coluzzii* mosquitoes swarming above a visual marker under simulated sunset conditions. We found that individuals alternate between straight flight and rapid turning maneuvers known as saccades. These saccades are triggered at the edge of the swarm, and their directions are biased along the sunset direction. This behavior was found consistent between both datasets and across swarm sizes, including solitary individuals, indicating that inter-individual interactions play a limited role within the studied size range.

We developed a simple agent-based model incorporating three behavioral rules: attraction to the swarm center, alignment normal to the sunset horizon, both driven by environmental cues, and repulsion through short-range collision avoidance. The model reproduced key features of natural mosquito swarms, including looping flight paths, central density peaks, and directional alignment. Even in the absence of direct inter-individual interactions, the model generated realistic emergent swarm dynamics, demonstrating that social coordination is not required for swarm emergence.

Our findings suggest that mosquito swarming is primarily guided by environmental perception rather than social interaction. This minimal framework offers a new perspective on insect swarming and may apply broadly to the many other insect species that form mating swarms. Understanding these rules could inform strategies for vector control and improve the effectiveness of interventions targeting mosquito reproduction.

**Author Summary:** Mosquito swarms are a familiar sight in many parts of the world, but the rules that govern their movement have remained mysterious. In our study, we investigated how male malaria mosquitoes behave in mating swarms. Using high-speed videography and computer simulations, we discovered that these swarms do not rely on complex social interactions. Instead, each mosquito follows a few simple rules based on visual cues from the environment, like the position of the sunset and a ground marker. These rules are enough to produce the looping flight patterns and dense swarms observed in nature. Surprisingly, even a single mosquito can perform this behavior in isolation.

This minimal approach challenges the idea that swarming requires strong social coordination. It also opens new possibilities for controlling malaria mosquito populations by targeting the environmental factors that influence mating behavior. Since many insects rely on swarming to reproduce, our findings may help explain similar behaviors across species and inform broader strategies for insect population management.

## Introduction

Few behaviors are as mesmerizing as the coordinated movements of flocking, schooling or swarming animals. Such collective behaviors can serve the purpose of foraging, mating, migrating, or protecting against predators [1–6]. Studying swarms offers insights into social learning, information transfer and animal ecology [7,8]. One key discovery about these apparently complex behaviors is that they seem to emerge from simple behavioral rules [9,10]. The “boids” model suggests that animals follow three rules based on distance: attraction to distant individuals, alignment with nearby neighbors, and repulsion to avoid collisions. Our understanding of bird flocks and fish schools is largely based on these rules and models that integrate them [11–15], but it is still unknown if flying insects in swarms follow these rules or how these swarming insects achieve coordination [16–18].

Insects exhibit a wide range of collective behaviors, from coordinated migratory swarms of locusts [19], to task-oriented aggregations of eusocial species such as honeybees [20], and the mating swarms of mosquitoes and midges [16,21–23]. Unlike the strong alignment seen in vertebrate groups [24–27], many insect swarms appear comparatively chaotic [16,18,21,24–31].

Mating swarms of mosquitoes are temporary aggregations formed by males near visual markers at dusk or dawn, where females visit to find mates [28,29]. These swarms act as leks, spatially fixed arenas for sexual selection, rather than migratory collectives. Extensive research has characterized their spatial structure and environmental triggers [16,21–23,30,31], yet whether individuals interact strongly with conspecifics remains debated [16,18,23,32]. Some studies report flight-path correlations suggesting attraction or alignment [17,23,32], while others attribute these patterns to simultaneous responses to external cues such as swarm markers and light gradients [21,30,31]. Quantitative models are needed to resolve these uncertainties [16,21–23,32–36,36–41].

Mating swarms have been described to exhibit looping flight paths [21,22,28,33]. This led to the modeling of mosquito swarming as damped harmonic oscillator coupled to the swarm centroid [23,33], and to the claim that male mosquitoes follow the boids model behavioral rules [17]. The proposed behavioral rules are based on the assumption that individual mosquitoes are attracted to the perceived center of the swarms, and that swarming mosquitoes align their flight direction with those of their neighbors. However, while there are quantitative evidences of collision avoidances between mosquitoes [18,23], mutual attraction and alignment remain unproven.

The argued attraction and alignment among swarming mosquitoes are based on observed correlations in their flight paths [17,23]. However, these correlations might instead arise from simultaneous responses to environmental cues like a swarm marker and the sunset [21]. This is further supported by the importance of environmental cues for swarming of many insects [34], and the limited sensory capabilities of male mosquitoes. Most mosquitoes swarm in low light conditions at dusk, and their hearing is tuned to detect female conspecifics rather than other swarming males [35]. Thus, given that male mosquitoes likely cannot accurately detect other males beyond a few centimeters [18,36], their capacity for mutual attraction or alignment is limited.

Here, we aim to identify the behavioral rules that govern swarming in male *Anopheles coluzzii* mosquitoes. Building on our previous work that revealed the environmental cues enabling stable swarm formation [32], we combine detailed swarm kinematics analyses with agent-based modeling to test whether complex swarm dynamics can emerge from simple, minimally interactive rules. We base our swarm-kinematics analyses on two published datasets containing three-dimensional flight tracks of individual swarming mosquitoes [32,37], and use the empirically derived kinematic patterns to parameterize and test a minimal agent-based model.

Specifically, we investigate the relative roles of environmental cues and conspecific interactions in shaping swarm organization. To address this, we adopt a bottom-up, mechanistic approach: (1) use the published swarming datasets to analyze and characterize the three-dimensional flight kinematics of individual mosquitoes and the emergent swarm structure [32,37]; (2) infer candidate behavioral rules from these data and prior literature [18,31,32,37,38]; (3) implement these rules in a physics-based agent-based model; and (4) evaluate whether this minimal rule set reproduces the experimentally observed swarm-level dynamics. This integrated framework allows us to directly compare empirical observations from literature with model predictions and assess the necessity of social interactions for swarm cohesion.

## Materials and Methods

In this study, we combine analyses of published three-dimensional swarming datasets with agent-based modeling to identify the behavioral rules that generate mosquito mating swarms. Our workflow consists of two components: (i) extracting and characterizing the flight kinematics and emergent swarm structure from the existing empirical datasets [32,37], and (ii) implementing these empirically derived patterns in a computational agent-based model to test whether simple rules are sufficient to reproduce observed swarm dynamics.

For our swarm-kinematic analyses, we used two empirical datasets published by Poda et al. 2024 [32], and Feugère et al. 2022 [37], hereafter referred to as the *Poda dataset* and the *Feugère dataset*, respectively. Specifically, we use the Poda dataset as the primary empirical reference and the Feugère dataset for generalization. Both datasets contain reconstructed three-dimensional trajectories of individual swarming mosquitoes. We used both datasets to identify and characterize the flight patterns of individual swarming mosquitoes and the emergent structure of the swarm.

We then used our primary high-fidelity Poda dataset to develop and parameterize the agent-based model; this dataset provides the most continuous and detailed swarming trajectories under conditions closely aligned with our modeling assumptions (e.g. being above of a flat and square swarm marker). Finally, we evaluated the model by comparing its output to the kinematics observed in both datasets, allowing us to assess how well the modelled behavioral rule set captures empirical swarming behavior and to test its generality across the two independent experimental conditions.

### Experimental animals

The empirical datasets analyzed in this study were generated using laboratory-reared male *Anopheles coluzzii* mosquitoes, as described in Poda et al. 2024 [32] and Feugère et al. 2022 [37]. In both studies, mosquitoes originated from recently established laboratory lines derived from wild females, were reared under controlled insectary conditions, and were 4–6-day-old at the time of swarming recordings. The Poda dataset contains swarming recordings of males only, whereas the Feugère dataset includes separate recordings of male and female swarms. Further details on rearing conditions, environmental parameters, and experimental procedures are provided in the original publications [32,37].

### Experimental setup

The *Poda* and *Feugère* datasets were both generated in controlled laboratory environments, in flight arenas equipped with a black ground marker to induce swarming under simulated sunset illumination. The most relevant experiment-specific details are summarized below.

In the *Poda dataset* [32], swarms were recorded in a transparent plexiglass flight arena measuring 2.0 × 0.7 × 1.8 m and placed within a larger experimental room (5.1 × 4.7 × 3.0 m). A 40 × 40 cm black cloth on the arena floor acted as the swarm marker. Sunset was simulated by gradually dimming overhead lights while maintaining a dedicated sunset light behind a black horizon panel. Two side-mounted near-infrared cameras (Basler acA2040-90umNIR) equipped with IR band-pass filters recorded at spatial and temporal resolutions of 2048 × 2048 pixels and 50 fps, respectively; four near-infrared illuminators provided minimally-invasive videography illumination.

In the *Feugère dataset* [37], swarming was recorded in a larger and more open flight arena, thus reducing potential wall-interaction effects. Illumination differed from the Poda setup: infrared sources were placed above and behind the swarm marker rather than laterally. The swarm marker itself also differed in geometry, consisting of a 30-cm-diameter slanted vertical cylinder instead of a flat square marker. Acoustic stimuli were delivered during recordings, although the original study showed little effect of this on overall swarm dynamics [37]. Three-dimensional flight tracks were reconstructed at the same 50 Hz recording framerate, allowing the same kinematic-analysis pipeline to be applied as for the Poda dataset.

### Experimental conditions

In the *Poda dataset* [32], swarming behavior was recorded across six laboratory swarming events, each consisting of three 5-minute recording periods corresponding to the onset, peak, and end of swarming activity. For each event, 30 or 50 male *An. coluzzii* mosquitoes were introduced into the flight arena one hour before dusk to allow acclimatization. This produced swarms ranging from a single initiator to up to 26 simultaneously swarming males, enabling analysis of swarm-size effects within the experimentally observed range. Swarming began when the overhead lights were fully dimmed and only the simulated sunset light remained illuminated.

In the *Feugère dataset* [37], swarming recordings were obtained under comparable dusk-transition light conditions, using the same dusk-onset timing to induce swarming. Separate experiments were conducted for male and female swarms, providing independent datasets for both sexes. As in the Poda study, swarms were recorded for several minutes once stable swarming was established. Acoustic stimuli were presented during recordings, although analyses in the original publication showed minimal influence of these stimuli on overall swarm dynamics [37].

### Flight kinematics analyses

The swarming flight path reconstruction resulted in three large database of flight tracks, being the Poda dataset and the Feugère *male* and *female* datasets. Each dataset consisted of a series of flight tracks, each defined using a space-time array (**x, y, z, t**), where **t** is the time stamp array throughout the flight track, and (**x**,**y**,**z**) is the position array of the mosquito relative to the center of the swarm marker ((*x,y,z*) = (0,0,0)). The *x, y* and *z*-coordinate axes are directed parallel to the sunset horizon, normal to the sunset horizon, and vertical up. For convenience, the *z*-coordinate is generally referred to as height, and the *y*-axis as the sunset axis (pointing towards the sunset).

We further analyzed these flight kinematics data using Matlab 2022b (MathWorks). First, we smoothed the track positions using a Savitzky-Golay filter. Then, we estimated for each flight track the velocity arrays (*u, v, w*) and acceleration array (*a_x_, a_y_, a_z_*), using a central finite difference scheme of second-order accuracy. From the velocity array, we estimated the angular speed throughout each flight track as in [39]. Finally, we determined the temporal dynamics of angle of view of the visual swarm marker along *y*-axis (α) [32].

Diptera commonly fly in relatively straight flights, interspersed with rapid turning maneuvers, commonly called body saccades [40,41]. To test whether our swarming mosquitoes also do so, we used an analysis approach inspired by the study of van Breugel et al. (2022) on saccadic flight maneuvers in fruit flies [42]. To quantify when, where and how often mosquitoes were flying straight or performing rapid turning maneuvers, we identified where their angular speeds were close to zero (= straight flight) and where it peaked (= turning maneuver). For that, the temporal positions of peaks and valleys (i.e. local maxima and minima, respectively) of angular speed were detected using the Matlab function findpeaks, with a minimum prominence of 300° s^−1^ (Fig. 1C). This threshold was the same as the one used in the previous Diptera flight study [42]. Furthermore, we performed a sensitivity analysis in which we systematically varied the threshold value (Fig. S1). This showed that the chosen threshold allows for a good separation between turning maneuvers and straight flight paths, and that the set threshold value has little impact on the main results (Fig. S1).

**Fig. 1.**
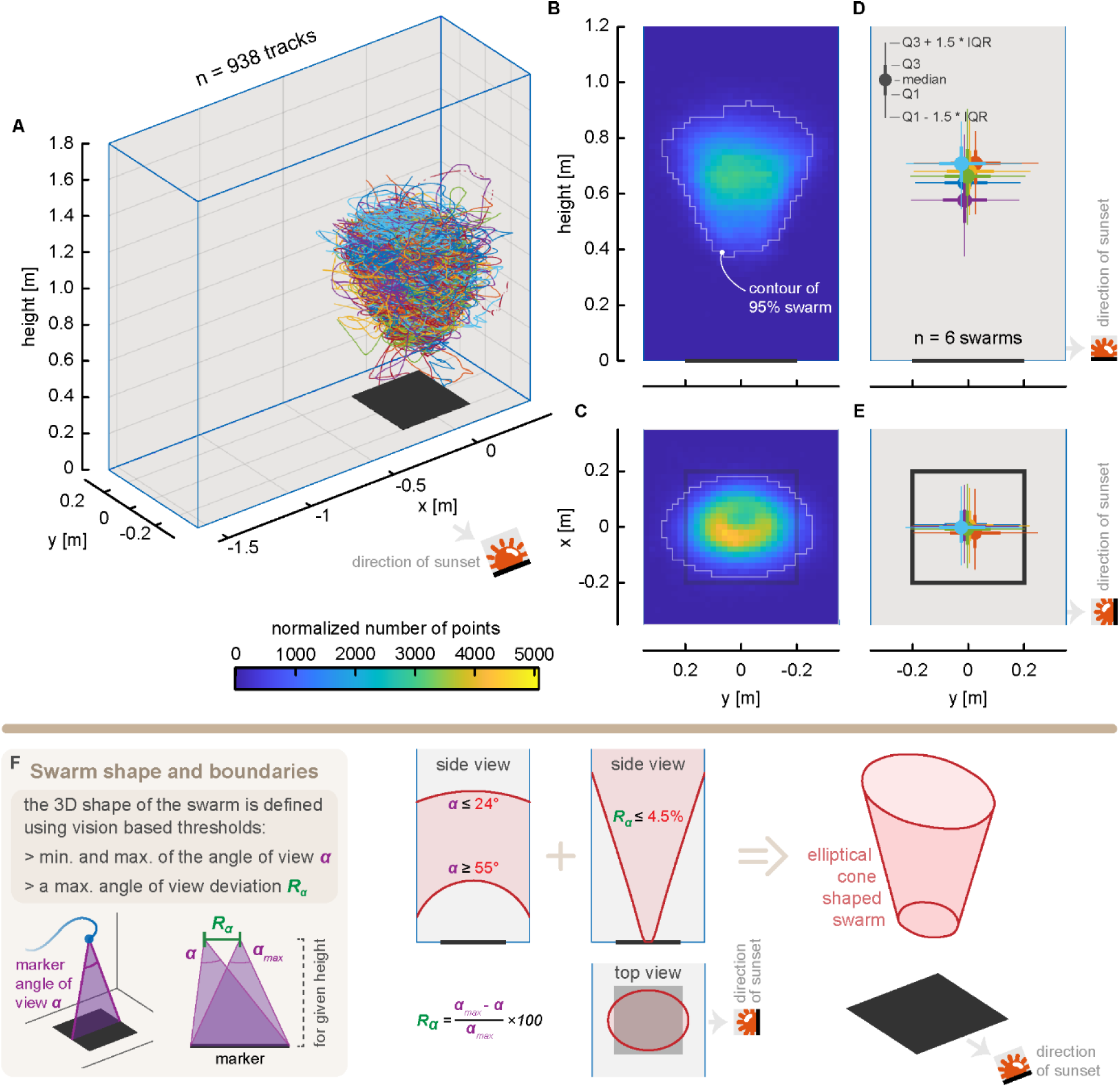
The characteristics of male *Anopheles coluzzii* mosquito swarms. (**A-E**) Experimental swarming data are from the Poda dataset [32]; see Fig S12 for equivalent data from the Feugère datasets [37]. (**A**) All 938 3D flight tracks of swarming male mosquitoes above a black ground marker, and inside the flight arena. (**B,C**) Side view (**B**) and top view (**C**) of the positional density distribution of mosquitoes flying within the swarm, defined as the total number of recorded mosquito positions per cell (2×2cm) for all replicates (*n*=6 swarms × 3 recordings per swarm = 18). The contour of the swarm comprising 95% of the data points is shown in white. The swarm marker is shown as a black line or square, and the sunset direction is shown on the right. (**D,E**) Side view (**D**) and top view (**E**) of the median location and quartiles of each of the six recorded swarms. (**F**) Legend and schematics explaining how the three-dimensional shape of the swarm above the square marker is defined using three thresholds of the marker’s visual angle α (from [32]).

The temporal positions of angular speed peaks and valleys were then used to divide each flight track into straight flight sections and saccadic turns. For each straight flight section between two saccadic turns, we computed the following flight kinematics parameters: flight distance travelled *d*, Euclidian distances *d*_E_, flight path tortuosity (τ=*d*/*d*_E_), duration (Δ*t*), flight speed (*U*), climb angle (γ), and flight heading relative to sunset axis (azimuth angle ψ=0° defines flight towards the sunset) (Fig. 1D-I). For each saccadic turning maneuver, we estimated the turn angle (Ω) as the difference in flight direction between the straight flight paths before and after the turn (Fig. 1D,G). For the Poda dataset, we determined the distance to the nearest walls (*d*_wall,*x*_ and *d*_wall,*y*_), to test for potential wall effects. We used these metrics to characterize the individual flights of swarming mosquitoes, and test the effect of environmental cues and those of conspecifics on the swarming flight kinematics.

### Spatial distribution of swarming behavior

To determine the spatial distribution of the flight kinematics within the swarm, we computed a series of kinematic parameters and visualized them in combined side-view and top-view projections [39]. For this, the experimental volume was divided in smaller sub-volumes of cross-section equal to 2×2 cm (Fig. 2). In each sub-volume, we calculated the relevant flight kinematics parameters and visualized the results within the two projections: a side view and a top view. For these visualizations, we used heatmaps for scalar parameters and vector fields for vectorial parameters.

**Fig. 2.**
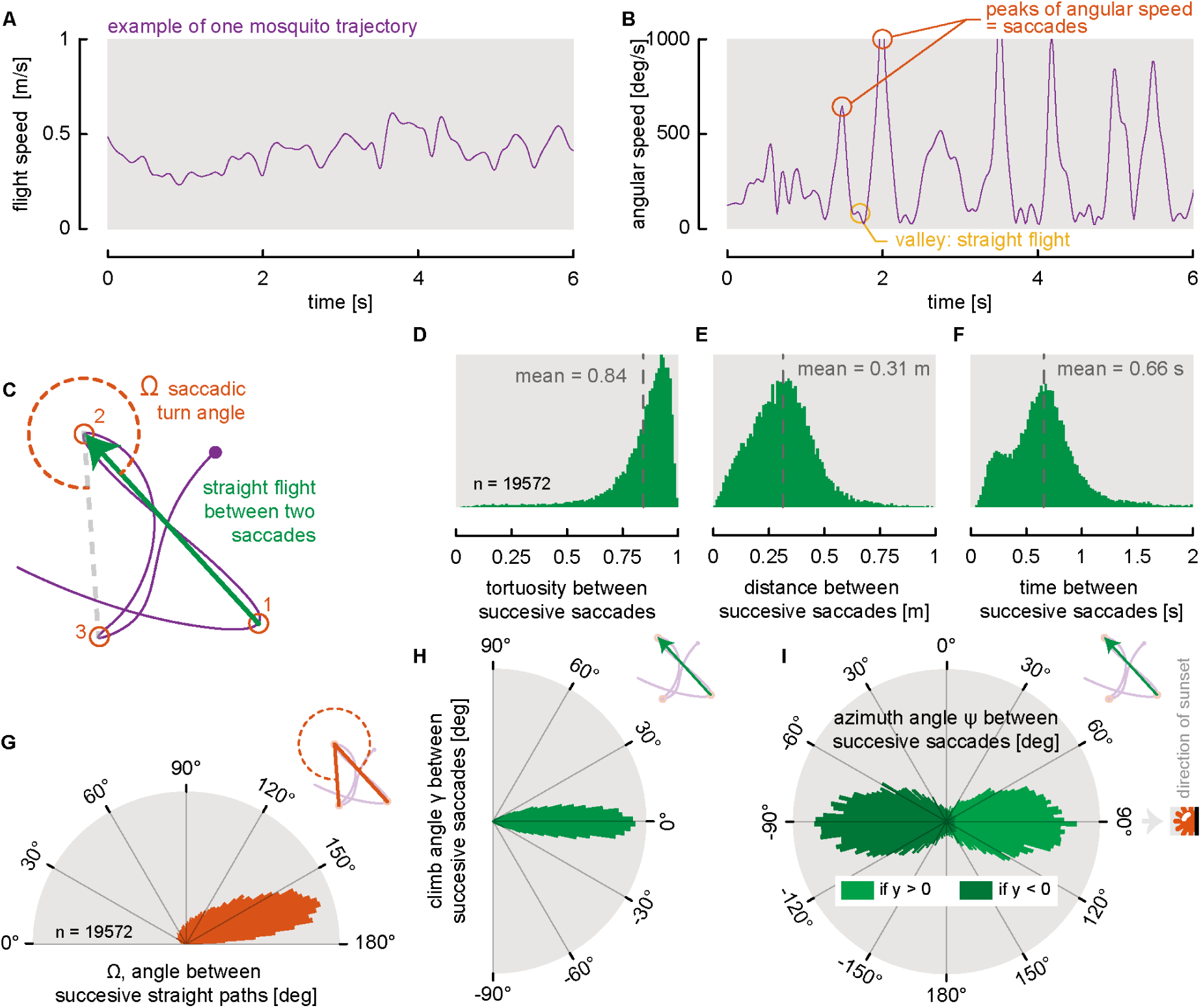
Swarming flight tracks consist of series of straight flight paths and saccade turns. Experimental swarming data are from the Poda dataset [32]. see Fig S13 for equivalent data from the Feugère datasets [37]. (**A,B**) Temporal dynamics of flight speed (**A**) and angular speed (**B**) of an example swarming flight trajectory. These metrics are oscillating over time in a seemingly periodic way. Local maximums and minimums (i.e. peaks and valleys) of angular speed are identified to highlight the part of the track when the mosquito was flying straight (angular speeds close to zero), or exhibiting rapid saccadic turning maneuvers (high angular speeds). (**C**) Top view of a swarming flight trajectory, including three consecutive saccadic turns (1,2,3; defined in orange), the straight flight path between saccades (green vector) and the saccadic turn angle (Ω). (**D**,**E,F**) Histograms showing the tortuosity (**D**), distance traveled (**E**), and time duration (**F**) of the straight flight phases between successive saccadic turns (*n*=19,572 straight flight phases within 938 trajectories). (**G**) Polar histogram of the saccadic turn angle Ω (*n*=19,572 saccades). (**H,I**) Polar histograms of the climb angle (i.e. angle from the horizontal plane; **H**) and azimuth angle (i.e. angle on horizontal plane; **I**) during the straight flight phases. Azimuth angles of −90° and +90° correspond to directions away and towards the sunset, respectively (see sunset symbol in **I**).

The scalar flight kinematics parameters of which we estimated the spatial distribution are: flight position density, linear and angular flight speed, saccade-to-straight flight ratio, straight flight path length, straight flight duration, straight flight heading (azimuth angle ψ), straight flight climb angle γ, saccadic turn angle Ω, and distance to the closest neighbor. The vectorial flight kinematics parameters were the local mean flight acceleration.

The spatial distribution of flight position density was computed as in Poda et al [32]. These mean track densities were normalized by the maximum number of data point in one sub-volume. To visualize the acceleration vector field, we computed the mean acceleration components within each sub-volume and used these to draw a single representative vector in the *xy*-plane (top view) or *yz*-plane (side view).

#### Agent-based modelling

To test which behavioral rules are needed to reproduce the swarming dynamics observed in the Poda and Feugère datasets, we developed a minimal agent-based model informed by the empirical flight kinematics extracted from these datasets. The model treats mosquitoes as self-propelled agents whose motion is governed by a small set of behavioral rules derived from measured straight-flight and saccadic-turn statistics. Below, we describe how empirical kinematic distributions were used as model inputs, how the behavioral rules were formulated, and how the resulting *in silico* swarms were simulated and analyzed.

### Experimental data as input for the agent-based model

We used the Poda dataset to estimate the distributions of climb and azimuth angles of the straight flight paths between two saccadic turns, relative to the position of the preceding turn (Fig. S2). We visualized these correlations using vector fields (Fig. S2*a,g*) and as probability density distributions of flight heading and climb angle versus angular position of the turn (Fig. S2*b,h*). We then fitted linear trends to these density distributions (Fig. S2*c,i*) and computed the distributions of flight directions around these linear fits (Fig. S2*d,j*). These combined linear-trend functions and residual distributions were used as direct inputs for the agent-based model. In simulations, after each saccade, the new flight direction was drawn by first evaluating the linear trend at the saccade location and then sampling from the corresponding residual distribution (Fig. S2e,k), which accurately reproduced the empirical post-turn directional distributions (Fig. S2f,l).

This heading and climb-angle relationship is the only component of the model directly fitted to the Poda dataset; all other parameters were taken from previous empirical observations in literature and were not tuned to improve agreement with the empirical data. This ensured that the correspondence between simulated and observed swarm dynamics arises from the minimal behavioral rule set rather than parameter optimization.

### Identifying agent-based model rules

By combining the results of our analyses of the Poda and Feugère datasets (Figs 1-3, S2, S12-14) with those of previous research [18,32,38,43], we identified candidate behavioral rules that may underly swarming of malaria mosquitoes. The three behavioral rules identified are:

**Fig. 3.**
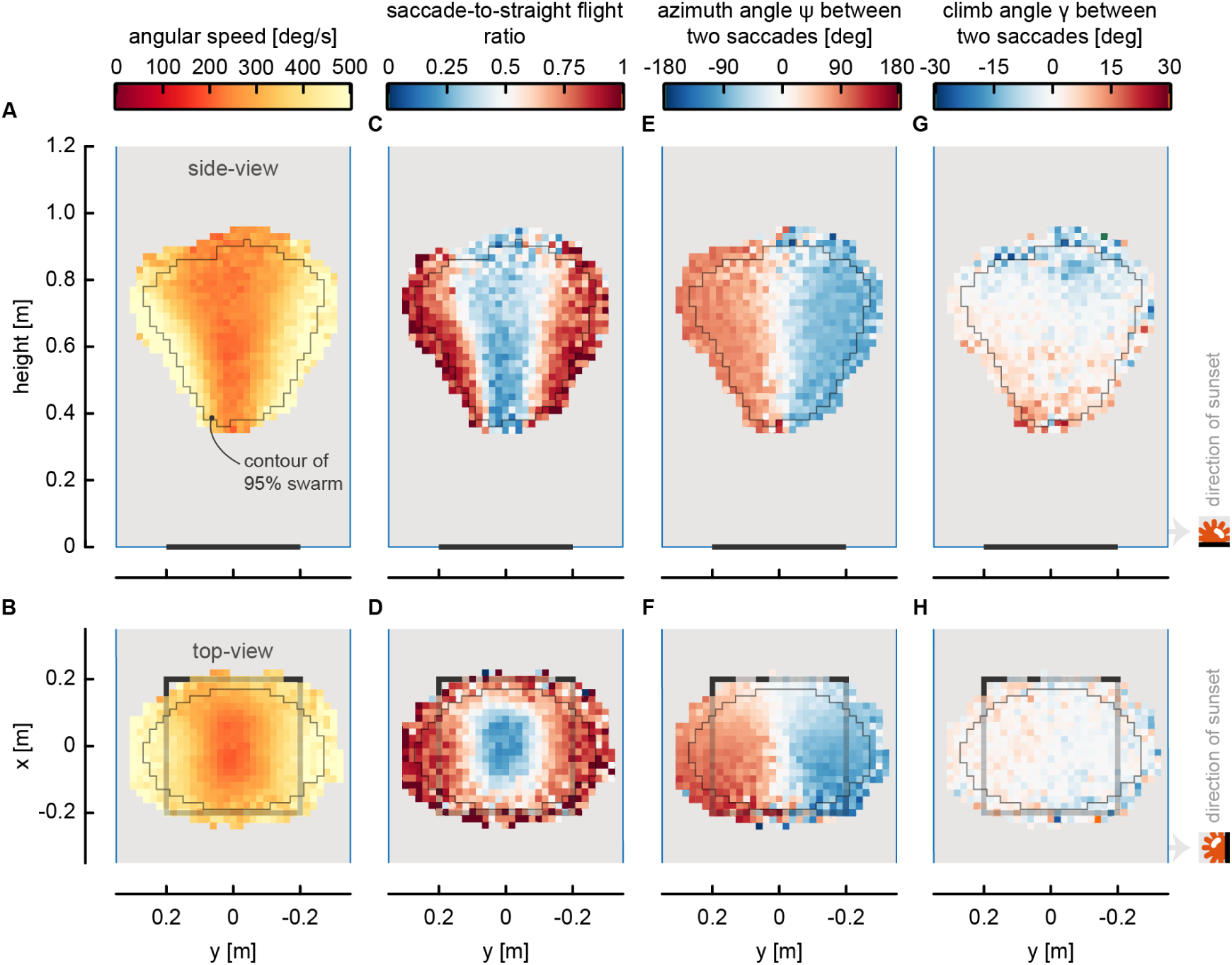
Spatial distributions flight kinematics parameters within the swarm. Experimental data are from the Poda dataset [32]; see Fig S15 for equivalent data from the Feugère datasets [37]. Side view (top row) and top view (bottom row) of the spatial distributions of the major flight kinematics parameters of swarming mosquitoes (*n* = 938 trajectories), including angular speed (**A,B**), the saccade-to-straight flight ratio (**C,D**), the straight flight azimuth angle (**E,F**), and the straight flight climb angle (**G,H**). The thin grey line shows the contour of the volume comprising 95% of the three-dimensional tracking data, the black bar and square represent the swarm marker, and the sunset location is indicated on the right. Data was shown at a cell size resolution of 2×2cm. No data was shown for cells with less than an average of one track per recording period and per swarm (i.e. 3 recording periods * 6 swarms = 18).

#### Behavioral rule 1

Swarming mosquitoes preferentially fly straight through the swarm center, where the marker viewing angle is maximal. They then continue flying straight, until the marker viewing angle falls below a threshold value, which triggers a turning maneuver [32].

#### Behavioral rule 2

The turning maneuver consists of a rapid saccadic turn within the horizontal plane, with biases towards the swarm center and along the sunset axis (towards and away from the sunset position) [32]. After the turn, the mosquito continues to fly in a straight line (rule 1).

#### Behavioral rule 3

Mosquitoes avoid collision when the distance to a conspecific decreases to within a few body lengths [18].

### Agent-based model development

To test how well this set of behavioral rules results in the observed complex emergent swarm dynamics, we developed an agent-based model in Matlab 2022b. The model is similar to the ones developed for flocking birds and schooling fish [11–18], but here agents primarily interact with environmental cues (the swarm marker and sunset position) rather than with conspecifics. Note that we do not aim to identify the exact sensory cues used by swarming mosquitoes, but focus on the response dynamics to positional and directional environmental cues. We structured the model description following the ODD (Overview, Design concepts, Details) protocol [44], to improve reproducibility and clarity.

Specific model parameters were taken from the Poda dataset and its accompanying publication [32]. We used the Poda and Feugère datasets together to validate the model and to assess the generality of the inferred rules across recording conditions. We applied the behavioral rules to the model using the following model rules (Fig 4):

**Fig. 4.**
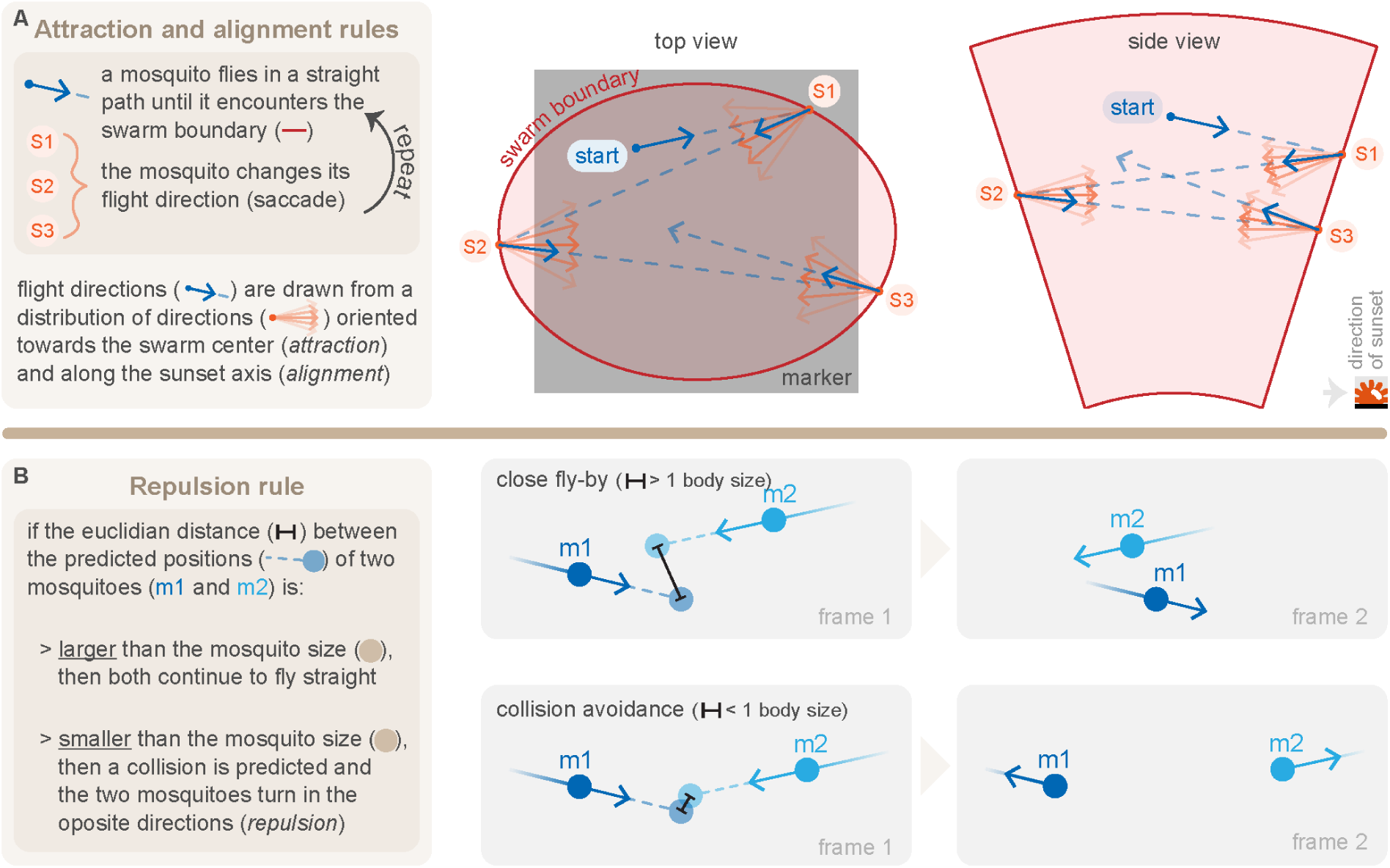
The behavioral rules used in our agent-based mosquito swarming model. (**A**) Legend and schematics describing the attraction and alignment behavioral rules of swarming male malaria mosquitoes. Swarming mosquitoes alternate between (i) straight flight across the center of the swarm volume and (ii) a saccadic turn at the swarm boundaries. During each saccade, they redirect their flight towards the swarm center (attraction) and along the sunset axis (alignment). (**B**) Legend and schematics defining the repulsion rule used in our model to prevent collision between close mosquitoes.

#### Model rule A - Initialization and process overview

A swarming mosquito starts at a random location above the marker, flying in a random horizontal direction at a constant speed of 0.5 m/s. This flight speed matches the mean reported in the Poda dataset [32].

#### Model rule B - Submodel: Saccadic turning

During this straight flight section, the simulated mosquito is constantly sampling the marker viewing angle, normal to the sunset horizon α (Fig. 1F). Based on this, the swarming mosquito performs a saccadic turn maneuver if (Fig 1F) [32,42]:

a. the marker viewing angle α becomes larger than 55° or smaller than 24°, i.e. when the mosquito reaches the bottom or top boundary of the swarm, respectively.
b. the marker viewing angle deviation *R*_α_ is larger than 4.5%. The marker viewing angle deviation is defined as *R*_α_= (α_max_-α)/α_max_·100%, where α is the current viewing angle and α_max_ is the maximal viewing angle for the current flight height (i.e. right above the marker). This lateral trigger corresponds to the elliptical cone-shaped sides of the swarm.

#### Model rule C - Stochasticity and submodel: Turn angle selection

The new flight azimuth and climb angles after each saccadic turn were randomly sampled from distributions based on the empirical trend-plus-residual distributions derived from the Poda dataset (Fig. S2).

#### Model rule D - Submodel: Collision avoidance

During straight flight, each mosquito constantly predicts the distance to its nearest neighbor *d*_predict_ at a future time step Δ*t*, using linear extrapolation from current position **x** and velocity **u** (**x**_predict_ = **x** + **u**·Δ*t*). We set the prediction time step equal to the simulation time step (Δ*t* = 0.02 s). If *d*_predict_ falls below one body length (∼6 mm), both mosquitoes execute a rapid evasive maneuver. Motivated by observed banked evasions [45], we implement this as an instantaneous 180° reorientation (velocity-vector inversion).

#### Model rule E - Process scheduling

After both the saccadic turn or the evasive maneuver, the simulated mosquito continued to fly in a straight line at a constant speed of 0.5 m/s, until the next saccadic turn or evasive maneuver was triggered.

### Agent-based model simulations

We used an agent-based model of swarming mosquitoes to simulate mosquito swarming. A complete list of model parameters and their default values is provided in Table S1 (Supplementary Material), using ODD terminology [44]. The *in-silico* swarm experiments were done in the same conditions as for our *in-vivo* experiments. All in-silico swarm simulations were performed under the same arena and marker dimensions as in the Poda dataset.

For all simulations, we simulated ten simultaneously swarming mosquitoes, matching the typical number of simultaneously swarming individuals observed in the Poda dataset [32]. We initiated flight of these ten mosquitoes at the same time, where each had a different random initial position and flight heading (model rule A). The simulations were run at a temporal resolution of 50 Hz, which is the same as the empirical video frame rate. At each simulation time step, we updated the position of mosquitoes based on their previous position, and current flight speed and direction. We continued the simulations for two minutes, resulting in 6000 sampling points per flight track. We replicated each *in-silico* experiment 20 times, resulting in 200 simulated flight tracks per condition.

Using this approach, we performed two sets of simulations. one in which all model rules were active, and one in which the collision-avoidance rule (Model rule D) was disabled. This allowed us to test for the relative importance of this repulsion rule in mosquito swarming.

#### Comparing the agent-based simulation output with the Poda dataset results

To evaluate how well the agent-based model reproduces the swarming kinematics observed in the Poda dataset, we compared model output directly to the empirical trajectories using a unified analysis pipeline. Both the empirical data and the simulated trajectories were processed with the same codebase, which included smoothing flight tracks with a Savitzky–Golay filter and computing velocities and accelerations via a second-order central finite difference scheme. All kinematic variables were derived using identical methods for both datasets, ensuring consistency in filtering and metric computation. The smoothing procedure attenuates peak accelerations, particularly during saccadic or evasive maneuvers involving rapid turning, which results in simulated turning dynamics that are more comparable to the empirical Poda trajectories.

Our primary validation step consisted of a detailed comparison between the simulated swarms and the Poda dataset, focusing on the spatial distribution of the main swarm-kinematic parameters. We visualized the in-silico and Poda trajectories using combined top- and side-view projections, employing heatmaps for scalar parameters and vector fields for vectorial parameters, as described above.

A second test of model fidelity involved comparing heading and climb-angle behavior between the simulations and the Poda dataset, based on the dependence of straight-flight headings on azimuthal position within the swarm and the dependence of climb angle on flight height (Fig. S2). Because the saccadic-turn rule (Model rule C) was parameterized from the Poda dataset, these comparisons test whether the implemented distributions correctly reproduce the empirical relationships (experimental results in Fig. S2a–d,g–j; simulation results in Fig. S2e,f,k,l).

Next, we compared the temporal dynamics of speed and acceleration along straight-flight segments between successive saccades, for both the Poda dataset and the agent-based model simulations. This analysis tests whether the model captures the characteristic temporal modulation of speed and acceleration observed in empirical straight-flight segments.

Finally, we compared simulations with and without the collision-avoidance rule (Model rule D) to the Poda dataset, assessing whether short-range repulsion improves agreement with empirical nearest-neighbor dynamics (Fig. 5I–L). For both simulation variants, and for the Poda dataset, we estimated the nearest-neighbor distance and the acceleration directed toward the nearest neighbor at each time step. We then evaluated how this acceleration depends on neighbor distance, testing the hypothesis that collision-avoidance behavior produces negative (repulsive) accelerations at very short range [18].

**Fig. 5.**
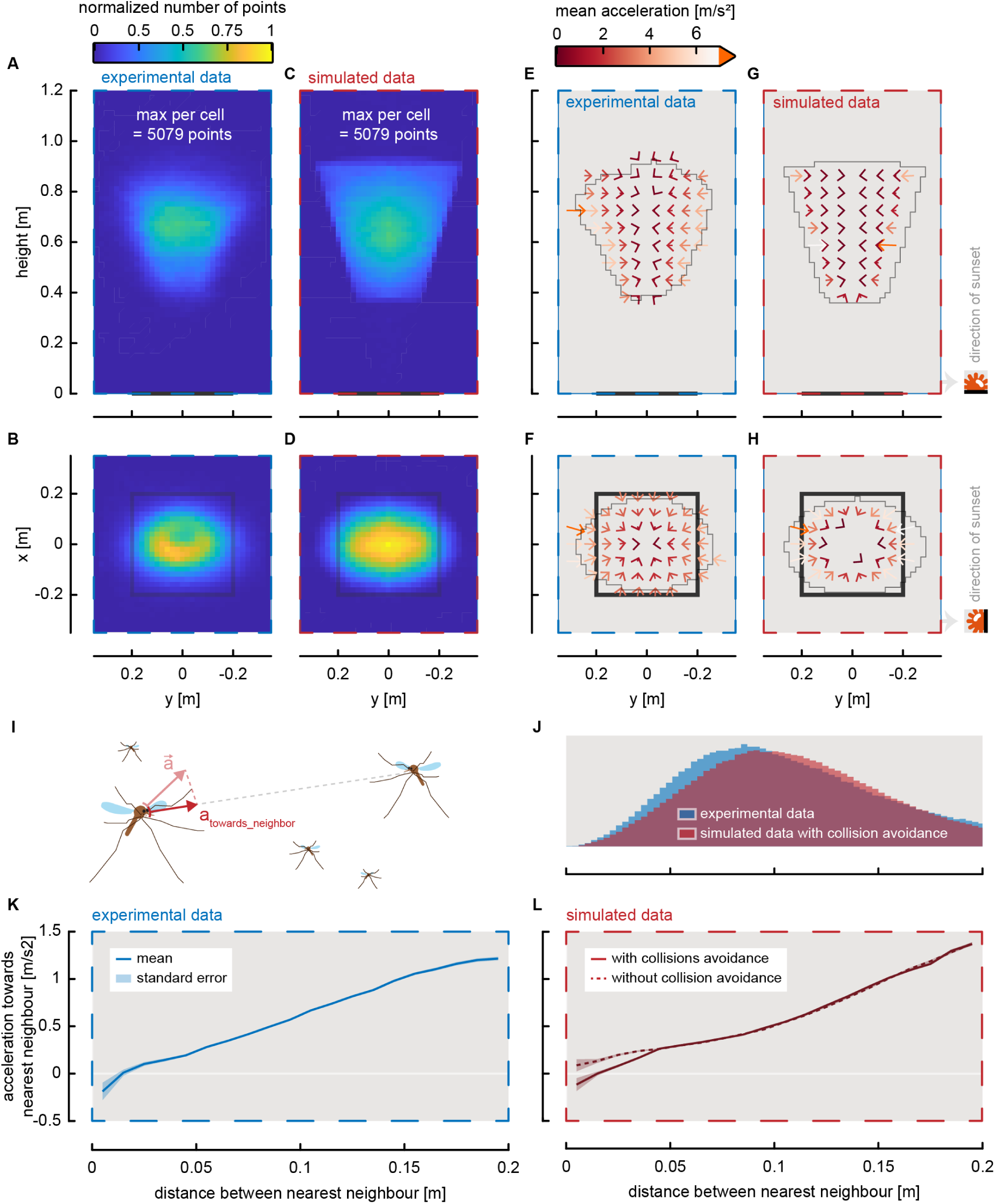
Comparison of the spatial characteristics within the experimental swarms and simulated swarms from the Poda dataset. Throughout, empirical results (*n*=938 tracks) and simulated results (*n*=938 tracks) are highlighted by blue and red dashed boxes, respectively. (**A***-***D**) Side view (**A,C**) and top view (**B,D**) of the positional density distributions, within experimental swarms (**A,B**) and simulated swarms (**C,D**). The simulated data was generated using the behavioral rules described in figure 4. (**E-H**) Side view (**E,G**) and top view (**F,H**) of the acceleration vector fields, within experimental swarms (**E,F**) and simulated swarms (**G,H**). The thin grey line shows the contour of the volume comprising 95% of the three-dimensional tracking data, the black bar and square represent the swarm marker, and the sunset location is indicated on the right. Data was shown at a cell size resolution of 2×2cm. No data was shown for cells with less than an average of one track per recording period and per swarm (i.e. 3 recording periods * 6 swarms = 18). (**I**) Schematic for defining the acceleration towards the nearest neighbor as the component of the acceleration vector of a mosquito directed towards its closets neighbor. (**J**) Histogram of distances between nearest neighbors for the experimental and simulated swarms. (**K,L**) The accelerations towards their nearest neighbors versus the distance between the two neighbors, for the empirical Poda dataset (**K**) and the simulated swarm (**L**). (**K**) In the experimental swarms, neighboring mosquitoes tended to accelerate away from each other (negative accelerations) only at very close distances (< 1.4cm), most likely to avoid collision. (**L**) Adding a simple collision avoidance mechanism to our simulations produced very similar negative accelerations between conspecifics at close distances (< 1.4cm). Without such collision avoidance mechanisms, no negative accelerations between neighbors were produced. (**K,L**) At larger distances, neighboring mosquitoes tended to accelerate towards each other for both the experiments and simulations, although no mutual attractions were modelled. This shows that highly correlated swarming behavior can emerge from responses to environmental cues only, and thus no mutual attractions and alignments between conspecifics are needed.

In addition to the detailed comparisons performed with the Poda dataset, we carried out a general, lower-resolution comparison with the Feugère dataset to assess whether the broad spatial and directional patterns produced by the model are consistent across independent experimental conditions. This comparison was not used for parameter fitting but served solely to evaluate the generality of the inferred behavioral rules.

## Results

### Empirical swarming results

We analyzed two published datasets containing three-dimensional trajectories of *Anopheles coluzzii* swarms [32,37]. The Poda dataset serves as the primary empirical reference, while the Feugère dataset is used to assess the generality of the observed spatial and kinematic patterns.

In the Poda dataset, male mosquitoes swarmed above a black visual ground marker under simulated sunset illumination, yielding 938 reconstructed 3D flight tracks from 18 recordings across six swarming events that spanned the onset, peak, and end of swarming (1–26 individuals; Fig. 1A). Consistent with previous analysis [32], these swarms displayed a characteristic upside-down elliptical cone shape above the marker (Fig. 1A,B). The major axis of the ellipse was oriented normal to the sunset horizon, and the minor axis was oriented parallel to it. Swarm density peaked at the center and decreased gradually toward the boundaries (Fig. 1B,C).

Following Poda et al. [32], the boundaries of the swarm volume can be described using the marker viewing angle α and its relative deviation from the local maximum value Rα. Mosquitoes remained within the swarm height range when 24° ≤ α ≤ 55°, and within the lateral boundaries when Rα ≤ 4.5% (Fig. 1F). Together, these criteria delineate the three-dimensional region in which swarming males are most likely to occur.

The Feugère dataset includes swarms of both male and female *An. coluzzii*, recorded under a different arena geometry and using a cylindrical ground marker. Despite these differences, the Feugère data exhibit similar large-scale swarm structure, including an elevated elliptical swarm volume and a central density peak (Fig. S12a–d). These similarities indicate that the primary geometric features of laboratory mating swarms are robust across experimental setups and do not depend on specific marker shape or arena configuration.

### Flight kinematics of individual swarming mosquitoes

In our previous analysis of the Poda dataset [32], we focused primarily on swarm-level structure and temporal dynamics. Here, we extend that work by quantifying the flight behavior of individual mosquitoes within the swarm (Videos S1–S4). Flight-track durations ranged from less than one second to 232 s, with an average of 15 s [32]. Swarming males exhibited stereotypic flight behavior, as expressed by the temporal dynamics of their linear and angular flight speeds (typical example in Fig. 2A,B). They alternated rapidly between two flight phases: relatively straight flight paths and rapid sharp turns. These rapid turns are often referred to as saccadic maneuvers, and are typically exhibited by flying Diptera [40,41].

We identified straight-flight segments and saccadic turns using the local minima and maxima of angular speed, respectively (Fig. 2B). On average, swarming mosquitoes exhibited saccadic maneuvers every 0.66±0.32 s (mean±standard deviation) (Fig. 2F). The resulting saccadic turn frequency of 1.5Hz is similar to the turn frequencies found in previous research on swarming mosquitoes [21,33]. During a single saccadic turn, swarming mosquitoes changed their flight direction with 165° (mode), and so they turned almost completely in the opposite direction. These turn angles during swarming are much larger than the typical turn angles of saccadic maneuvers previously described in foraging flies and host-seeking mosquitoes [40,46].

During the flight sections between consecutive saccadic maneuvers, swarming mosquitoes were approximately flying straight, as the tortuosity of these flight trajectories was close to one (τ=0.84±0.14; Fig. 2D). We therefore define these flight sections between saccadic maneuvers from here-on as straight flight sections. The distance traveled by mosquitoes in these sections was on average 31±14 cm, which is slightly less than the swarm marker size (of 40 cm) (Fig. 2E). These straight segments were primarily in the horizontal plane, and directed along the sunset axis (Fig. 2H,I; respectively). The climb angles γ during the straight flight sections were consistently close to zero (mode: γ=0°); Fig. 2H). To analyze the directionality of the straight flight sections, we determined the mean flight direction of each section, as defined by the azimuth angle ψ. These flight directions combined show a bimodal distribution (Fig. 2I): straight flight sections that followed a saccadic maneuver distally from the sunset were oriented primarily towards the sunset (*y*_saccade_>0: ψ=91.8° (mode); Fig. 2I), whereas mosquitoes flew mostly away from the sunset after a saccade proximally to the sunset position (*y*_saccade_<0: ψ=-91.8° (mode); Fig. 2I). Together, these patterns generated the characteristic back-and-forth looping flights aligned with the sunset axis.

Analysis of the Feugère dataset [37] showed similar individual-level kinematics for both male and female swarms (Fig. S13). Mosquitoes exhibited comparable saccade frequencies, turn angles, tortuosity values, and straight-flight climb angles (Fig. S13e,b,d). Straight-flight distances were shorter (Fig. S13c), likely due to the smaller cylindrical marker used in that study. As in the Poda dataset, straight-flight azimuths in the Feugère data showed a preference aligned with the sunset axis (Figs 2I and S13g, respectively). These similarities indicate that the observed kinematic patterns are robust across different experimental geometries and recording conditions.

### Variations in flight kinematics within the swarm volume

Next, we investigated how flight kinematics of the swarming mosquitoes varied with their position in the swarm. In the Poda dataset, we previously found that mosquitoes flew the fastest and accelerated the least in the swarm center [32]. In contrast, mosquitoes flew the slowest and accelerated most rapidly near the swarm boundaries, particularly in the regions adjacent to and distally from the sunset horizon [32]. We observed the same spatial trends in both datasets: the spatial distribution of angular speed within the swarm is very similar to the distribution of flight accelerations, whereby angular speeds were lowest near the vertical central axis of swarm, and highest on the swarm sides, adjacent to and distally from the sunset horizon (Poda dataset: Fig. 3A,B; Feugère datasets: Figs S14-15).

When looking at azimuth angles between successive saccades, the swarms can be divided in two halves: a region proximal to the sunset horizon, and one distally from it (Poda: Fig. 3E,F; Feugère: Figs S14–S15). After a saccade located near the sunset, mosquitoes primarily flew away from the sunset, and opposite. Similarly, mosquitoes near the top of the swarm tend to fly downward after a saccade, whereas near the bottom of the swarm, mosquitoes flew on average upwards after a saccade (Poda: Fig. 3G,H; Feugère: Figs S14–S15).

All together, these results shows that swarming mosquitoes, fly fast, horizontally and straight when crossing the swarm center, and perform rapid turns at the swarm boundaries (Poda: Fig. 3C,D; Feugère: Figs S14–S15). These kinematics are similar to those observed in swarming midges [43].

### The influence of experimental conditions on the swarming flight kinematics

In any behavioral experiment, it is important to test potential effects of the experimental conditions on results. Because the Poda dataset was recorded inside a plexiglass flight arena with limited dimensions, we quantified the distance between each saccadic turn and the nearest arena wall to assess whether turning events occurred close to the arena walls (Fig. S11). For turns at the swarm edge, distances to the side walls were 45 ± 8.5 cm and to the front/back walls 23 ± 7.1 cm, corresponding to ∼75 and ∼38 body lengths, respectively. These values are far greater than the short-range distances at which collision-avoidance behavior occurs (∼2.5 body lengths; Fig. 5K) [18], and exceed reported wall-interaction distances of one to a few body lengths [45,47].

As an additional check, we examined the Feugère dataset, which was recorded in a substantially larger and more open arena. In this dataset, the same turning patterns, boundary-localized saccades and elevated angular speeds at the swarm periphery, were observed (Figs. S14–S15), despite the much greater distances to any physical boundaries.

Together, the large wall distances in the Poda dataset and the recurrence of the same kinematic structures in the more open Feugère recordings provide indirect evidence that the spatial organization of turning behavior is not tightly coupled to arena proximity.

### Variations in flight kinematics with swarm size

In the Poda dataset, swarm sizes ranged from a single initiator male (Video S1) to groups of up to 26 simultaneously swarming individuals (Videos S2–S3). This range corresponds to the modal sizes reported for natural *Anopheles* mating swarms in the field, which commonly consist of several to a few dozen males [48–50], although much larger swarms, occasionally reaching hundreds to thousands of individuals, have been documented in some locations.

Within the Poda dataset, single initiators displayed the same looping straight-flight and saccadic-turn sequences as individuals in larger groups. To evaluate whether individual-level flight behavior varied over the swarm-size range present in the dataset, we quantified all saccadic-turn and straight-flight metrics for each swarm and compared them across group sizes (Fig. S8). No systematic differences were detected across swarm sizes, in saccadic turn angle, straight-flight distance, or straight-flight duration. Thus, within the small-to-medium swarm sizes captured in the Poda dataset individual kinematics were invariant with respect to group size.

### Behavioral rules of malaria mosquitoes swarming

The combined kinematic patterns observed in the Poda and Feugere datasets (Figs. 1–3, S8; S14-S15) show that male *An. coluzzii* follow a stereotyped sequence of straight-flight segments and rapid saccadic turns while swarming (Fig. 4; Videos S1–S4). Mosquitoes fly straight through the swarm center, where the visual angle of the ground marker is maximal (α_max_), and continue this straight flight until the viewing angle decreases below a threshold, at which point a turning maneuver is triggered (Fig. 1F) [32]. These turns are sharp saccadic rotations within the horizontal plane, biased toward the swarm center and along the sunset axis. Saccades also include vertical adjustments: individuals near the lower boundary of the swarm tend to turn upward, whereas those near the upper boundary tend to turn downward, consistent with the spatial pattern of marker viewing angles (in the Poda dataset: 24° and 55°, respectively). After each turn, mosquitoes resume straight flight until the visual-angle threshold is crossed again, initiating the next saccade. Previous work further shows that swarming mosquitoes perform collision-avoidance maneuvers at close range [18].

### Modelling and simulation results

Based on the Poda dataset results, we developed an agent-based model for simulating mosquito swarming. We used this model to test how well these simple behavioral rules can reproduce the complex emergent swarm dynamic observed in the Poda datasets (see Material and Methods). Agent-based models are powerful tools that simulate the interactions of individual agents to understand complex collective behaviors. By modeling each animal’s simple rules of movement and interactions, we can uncover how these lead to complex patterns, providing insights into social dynamics. The agent-based model reproduced key aspects of the kinematics observed in the Poda dataset, with simulated mosquitoes exhibiting swarm dynamics consistent with the empirical trajectories (Videos S1-S4, Figs. 5, S3-S7). Simulated mosquitoes primarily performed saccadic maneuvers at the swarm edges and flew mostly horizontally and straight through the swarm center, with flight directions oriented primarily along the sunset axis (Figs. S4 and S5). Still, these kinematics are directly resulting from the behavioral rules, and thus these strong similarities are expected.

More striking is the fact that the model generated emergent properties not directly tied to the specific rules. As in the Poda dataset, simulated mosquito density was highest in the swarm center (Fig. 5A-D), and distributions of straight flight path distances and durations were similarly distributed to those observed experimentally (Fig. S7). The model also replicated the radial acceleration field observed in real swarms, where mosquitoes accelerate towards the swarm center, with maximum acceleration at the edges and minimum at the center (Figs. 5E-H, and as in [31,51]). This field arises from saccadic turns at the swarm boundary followed by straight flights towards the vertical centerline.

Analysis of the Poda dataset revealed that neighboring mosquitoes accelerate away from each other at distances below ∼1.4 cm, likely due to evasive maneuvers [18]. In contrast, at larger distances, conspecifics tend to accelerate towards each other (Fig. 5K). These positive accelerations towards each other may be the by-product of simultaneous flight behaviors of conspecifics, not mutual attractions. We tested this in the model by comparing versions with and without a collision avoidance rule (Fig. 5L). Both models reproduced the positive acceleration towards neighbors at distances over 1.4 cm, suggesting that synchronized movements in swarms can emerge from reaction to environmental cues alone, without direct interactions between conspecifics (Fig. 5K,L). Still, only the model with collision avoidance captured the repulsive accelerations of real mosquitoes at close distances (Fig. 5K), showing that a minimal amount of interaction is needed to capture the emergent swarming behavior of mosquitoes.

A lower-resolution comparison with the Feugère dataset showed that the large-scale spatial and directional patterns generated by the model were consistent across datasets with different marker geometry and arena size (Figs 5 vs. S12–S15). This provides additional evidence that the modelled behavioral rules capture robust features of mosquito swarming.

## Discussion

### A minimal behavioral rule model for mosquito swarming

According to the principle of parsimony (or Occam’s razor), the hypothesis that requires the fewest assumptions, here minimally-interactive behavioral rules, should be tested before more complex hypothesis. This is what we did in our mosquito swarm study, where our combined analyses of empirical data from literature [32,37] with agent-based modelling show that the complex emergent swarming of malaria mosquitoes can be modelled based on a small set of minimally-interactive behavioral rules. Like in flocking bird and schooling fish models, these behavioral rules include attraction, alignment and repulsion rules. But strikingly, here the attraction and alignment behaviors are not towards conspecifics, but instead relative to external environmental cues: the swarming mosquitoes are attracted towards the region above the visual swarm marker, and they align their flight direction along the horizontal sunset axis (axis pointing towards the sunset position). In our simple agent-based model, the simulated mosquitoes used the angular view of the marker and the sunset direction as sensory cues for attraction and alignment, respectively. This resulted in very similar simulated flight dynamics than in real swarms (Fig. 5 and [16,22,23,32,37,38,43]).

In nature, mosquitoes might use additional environmental cues than in our model, or might even use different ones. For example, they might not or show different directional preferences in function of available environmental cues. Uncovering how exactly swarming mosquitoes use environmental sensory cues, such as the swarm marker and sunset light, will require experiments whereby the environmental cues are systematically manipulated, for example by moving the sunset point source or by varying marker shape and size. Still, our model shows that it is possible to generate swarming with realistic characteristics using simple minimally-interactive rules, and consequently the hypothesis that such minimally-interactive rules govern mating swarms should be preferred before assuming more complex rules.

Out of the three identified behavioral rules, we found that only the collision avoidance rule required to be interactive among conspecifics. Thus, collision avoidance can be regarded as a social response, but these occur very rarely (Fig. 5J), and only at very short distances between individuals (<2.5 body lengths; Fig. 5K) [18]. In contrast, birds and fish display high level of direct interactions, at comparable distances [26]. This shows that insect mating swarming might be inherently different from flocking and schooling, and that the previously suggested apparent long-distance interactions between swarming mosquitoes or midges [16,22,23,43] might in fact not be interactions, but instead the result of simultaneous responses to external environmental cues. This does not show that other types of interactions between swarming mosquitoes are not present, but it suggests that such potential interactions are not directly needed to produce stable mating swarms.

### Testing the minimal behavioral rule model

To test whether the identified behavioral rules capture the core dynamics of mosquito swarming, we developed an agent-based model implementing these rules, and compared simulation outputs (Fig.5) to those of our two empirical datasets (Poda dataset [32]: Figs 5, S3–S7; Feugère dataset [37]: Figs S12-S15). In the model, simulated mosquitoes fly along straight trajectories while continuously sampling the marker viewing angle α to assess whether they are about to cross the swarm boundary [32]. When a boundary-crossing is predicted, they execute a saccadic turn back toward the marker centerline, with new azimuth and climb angles drawn from kinematic distributions measured in the Poda dataset (Fig. S2). Thus, the model treats the swarm boundary as a binary threshold (inside vs. outside), producing sharply defined spatial zones where saccadic maneuvers occur versus straight-flight regions (Fig. 5C,D and S4c,d,g,h). In the empirical datasets, these zones are less sharply delineated (Figs 5A-B, S4a,b,e,f, and S12*a-d*), likely due to individual heterogeneity and perceptual noise in boundary estimation. For model parsimony, we did not incorporate such variability. Despite this minimalism, the model reproduces key empirical features: high mosquito density near the swarm center, acceleration vectors directed toward the swarm’s central axis, a concentration of saccades at the periphery and straight flights in the core, as well as a dominant flight orientation along the sunset axis (Figs 3-4, S3-S7, S13 and S15). Together, these parallels indicate that the simple interaction rules are sufficient to generate the principal characteristics of observed mosquito swarms.

Moreover, analysis of the Poda dataset showed that the swarming dynamics were largely invariant across swarm sizes, with the number of individuals (1–26), falling within the ecologically relevant range for *Anopheles* mating swarms [48–50], and having no measurable impact on key flight-kinematics metrics (Fig. S8). This confirms earlier findings that swarming remains highly stereotyped independent of group size [32], and supports the observation that even a single mosquito can display swarming behavior in the absence of conspecifics [21,52]. The low densities of mosquito swarms likely explain this invariance. Even in our largest groups, nearest-neighbor distances averaged ∼20 body lengths, far above the values typically observed in bird flocks (∼3–7 body lengths) or fish schools (∼1–2 body lengths) individuals rarely come close enough for neighbor-mediated interactions. Studies indicate that larger field swarms consist not only of more individuals, but are also of larger volumes [30,34,53], thereby maintaining similarly low densities. Together, these observations explain the lack of measurable density effects on swarm structure and kinematics, and suggest that potential interactions among males contribute little to the initiation or execution of saccadic turns under natural swarming conditions.

It is to be noted that even if current analyses show minimal male-male interactions, observed collision avoidances behavior may suggest that males exhibit territorial behaviors, typical in leks, potentially to increase their chance to encounter females and exhibit traits likely to be chosen by females [54,55]. In that case, some locations in the swarm might be more favorable for males. Such territorial effects would likely be easier to observe in very large swarms that can comprise thousands of mosquitoes [56], and thus would require dedicated studies. Similarly, further analysis of male-female interactions during swarming will be needed to better understand how the here-described behavioral rules affect male competitiveness and mating success.

### Conceptual and experimental limitations of our approach

In this study, we used a bottom-up mechanistic approach to test whether simple sensory–motor rules at the individual level are sufficient to generate the characteristic structure of mosquito swarms. This contrasts with top-down approaches that infer effective collective forces or statistical correlations from group-level data, including models of laboratory insect swarms that reproduce cohesion without specifying short-range behavioral rules [43,57]. Recent statistical-inference methods, such as that of Iacomelli et al. (2025), estimate interaction strengths from correlated trajectories in large laboratory swarms (>80 individuals) [58]. These correlation patterns suggest collective effects at high densities but do not resolve whether such effects are necessary for swarm formation more generally. Our bottom-up model is therefore complementary to these established approaches: statistical and effective-force frameworks quantify group-level patterns [43,57,58], whereas our mechanistic model tests how these patterns can emerge from explicit and minimal individual-level behavioral rules.[43,57,58]

We based our computational analysis on the Poda and Feugère datasets, both recorded under controlled laboratory conditions. Laboratory swarms inevitably differ from those observed in nature, but detailed three-dimensional kinematic recordings of natural swarms are not yet available. These two datasets therefore represent the highest-quality empirical material currently accessible, even though their recording environments differ from field conditions. The main differences relative to natural swarms are the following: (1) In both datasets, dusk conditions were created using artificial light sources, and the videography relied on infrared illumination. This differs from natural sunset light, which is parallel and partially polarized. (2) Swarming occurred in finite flight arenas. The Poda dataset was collected in a relatively small enclosed arena, whereas the Feugère dataset used a larger and more open space. As a consequence, potential wall-proximity effects are most relevant for the Poda recordings. (3) The number of released mosquitoes produced small to medium swarm sizes, ranging from single initiators to maximum group sizes of 26 and 6 individuals in the Poda and Feugère datasets, respectively.

We addressed these limitations as far as possible using the available data and within the scope of a computational study. Some limitations, however, can only be resolved experimentally, for example by manipulating wall positions or altering the properties and direction of sunset illumination. These experiments fall outside the present study but would be valuable for future work. Below, we discuss the main limitations of our modelling approach in more detail.

(1) In both the Poda and Feugère datasets, the preferred flight direction and the major axis of the swarm were along the sunset axis (*y*-axis), rather than with the geometry of the flight arenas. The recurrence of this directional structure across two independent datasets, recorded in arenas of different sizes and with different marker shapes, indicates that the alignment is robust to variation in the recording environment (Figs. 1-3, S12–S15). However, the present data cannot establish whether sunset cues directly cause the observed asymmetry in swarm shape. Demonstrating causality will require experiments in which the properties or position of the sunset cue are systematically altered. Our results do show that when a consistent external directional cue is present, whether sunset light or another environmental gradient, mosquitoes align their straight-flight segments along that axis. This alignment generates asymmetry in both individual flight dynamics and the emergent swarm structure.
(2) To assess whether the walls of the Poda arena influenced turning behavior, we quantified the distance to the nearest wall at the onset of each saccadic turn (Fig. S11). Saccades at the swarm boundary occurred at approximately 75 body lengths from the side walls and 38 body lengths from the front and back walls. These distances are far greater than the short-range interaction distances associated with collision-avoidance behavior (∼2.5 body lengths; Fig. 5K) [18], and exceed reported wall-interaction distances of one to a few body lengths [45,47]. As an independent comparison, the Feugère dataset, recorded in a much larger arena with a different marker geometry, showed similar swarm structure, turn dynamics, and directional tendencies (Figs. S12–S15). This cross-dataset consistency indicates that wall proximity is unlikely to be a major driver of the observed flight behavior. Natural swarms of anthropophilic *Anopheles* mosquitoes also often form near buildings or other vertical structures[56], so occasional proximity to walls is ecologically realistic. Taken together, these observations suggest that although wall effects cannot be entirely excluded, their influence on the behavioral patterns central to this study is likely limited.
(3) We compared the swarm sizes in the two laboratory datasets with values reported from natural *Anopheles* mating swarms [48–50]. The laboratory swarms fall within the ecologically relevant range, where field surveys typically report modal and median swarm sizes of a few dozen males. Within this size range, our analysis of the Poda dataset showed that swarm kinematics were largely invariant: key metrics such as straight-flight distance, turn angle, and turn frequency showed no measurable dependence on group size (Fig. S8). This agrees with earlier findings that swarming remains highly stereotyped across small to medium swarm sizes [32]. Field observations do, however, show a broad size distribution with a heavy upper tail, and swarms can occasionally reach very large sizes [48–50]. Our results should not be extrapolated to such extreme cases [58]. Future work that tracks individual flight trajectories in these larger, high-density swarms would be valuable for testing how well the minimal behavioral rules identified here generalize to denser and more populous swarms.

Overall, the swarm dynamics observed in our analyses are consistent with field descriptions of *Anopheles* mating swarms [53], but important limitations remain. Fully testing these limitations will require dedicated experiments, both in the laboratory and especially in semi-field and field settings. Such work should quantify and manipulate key environmental variables, including light polarization, environmental clutter, and swarm size and composition. These data would enable more precise evaluation of how environmental conditions shape individual behavioral rules and would allow refinement of the agent-based model. They would also permit tests of additional or alternative rules that may govern mating swarms under natural conditions. Finally, extending the model to incorporate variability in saccadic-turn dynamics and environmental noise will be important for assessing the robustness of the proposed behavioral framework in more realistic ecological contexts.

### Mating swarms in nature

It is plausible that many other insect species follow similar rules in mating swarms, although this requires further quantitative validation in a comparative framework (e.g. swaRmverse [59]). For example, *An. gambiae* mosquitoes have been observed to swarm near ground markers along an east-west axis [60], suggesting a comparable sunset-oriented alignment. In contrast, *Culex* mosquitoes display predominantly vertical movements during swarming [21], likely reflecting the influence of other environmental cues. Our approach could be used to systematically test such interspecific differences, particularly across the many species of swarming Diptera. It could also be used to explore how swarming adapts to changing environmental conditions such as wind or light perturbations [33,51,61], potentially revealing additional behavioral rules.

Our results do not address interactions between males and females within these mating swarms. Males are known to be strongly attracted to females entering the swarm [22,62], and it has been hypothesized that the high density of males in the swarm center allows them to quickly intercept females at the periphery [30]. Understanding the behavioral rules governing female interception is crucial for a complete understanding of dipteran mating swarms. Our finding that swarming males fly straight through the swarm center and perform sharp saccadic turns near its boundaries provides a mechanistic basis for how males may optimize encounters with incoming females. Moreover, the combined experimental and modeling framework developed here offers a powerful foundation for future investigations of male–female interactions within swarms.

Contrary to bird flocks and most fish schools, mosquito swarms act as leks in which males aggregate in a spatially fixed swarm that females visit to locate mates. These mating swarms also differ fundamentally from the collective behavior of eusocial insects such as honeybees, and from cooperative mass-migratory swarms seen in locusts [63,64]. In stationary mating swarms like those studied here, individuals are not required to achieve consensus decisions or cooperative task allocation as in eusocial groups, and their interactions can thus be governed by simple, local behavioral rules. Nevertheless, external perturbations, such as wind gusts or predator attacks, may still trigger coordination similar to the collective responses observed in social insects, birds, or fishes [15,16]. Importantly, eusocial insects represent only a tiny fraction of insect diversity across non-social taxa (e.g., Diptera, Ephemeroptera, Hymenoptera), likely exceeding the number of eusocial species by orders of magnitude [52]. Consequently, our results not only clarify the minimal interaction rules sufficient to generate mosquito swarming, but also provide a framework that could be applied to a large number of insect species that mate in swarms.

This understanding also has direct implications for disease vector control. By identifying the behavioral rules and flight dynamics that give rise to realistic swarming, our results provide a mechanistic basis for assessing how males locate, join, and maintain position within mating swarms. Such insights are particularly valuable for improving the quality and competitiveness of laboratory-reared males used in vector control programs. Reduced mating success of released males, whether sterile [62,65,66] or carrying gene drive [67,68], remains a major limitation to the efficacy of these approaches [69]. A deeper understanding of the flight characteristics, spatial positioning, and sensory cues that determine swarming behavior and mating success can guide the selection, rearing, or conditioning of males that perform more effectively in the field. More broadly, because mosquito swarms are spatially constrained, easily reproduced under laboratory conditions, and responsive to controlled perturbations, they represent a great experimental model for studying collective animal behavior in general and for translating such fundamental insights into applied solutions for vector-borne disease control.

## Acknowledgments

We thank our collaborator from the Human Frontier Science Program project for the numerous exciting discussions on mosquito swarming: Jeff Riffell, Ruth Müller, Saumya Gupta, Simon Sawadogo and Sofia Vielma. We thank Lionel Feugère for our scientific discussions. We thank Somda Stephane, Sougué Emmanuel, Guinko Nourou, Coulibaly Valérie and Bandaogo Abdoul Malik for mosquito rearing, and Charles Nignan and Domonbabele François de Sales Hien for assisting in the data collection. We thank Mathurin Fatou and Pie Müller for helping to install and calibrating the video tracking system.

## Author contributions

BSP, OR, AD and RKD conceived the original study, BSP and OR conceived the experimental design, and BSP, AC and FTM conceived the modelling approach. BSP performed data collection and the tracking analysis. AC performed data analyses, the modeling, and drafted the figures and manuscript. FTM, BSP and OR critically revised the manuscript. All authors revised the manuscript, gave final approval for publication and are accountable for the work performed therein.

## Data and materials availability

All original Matlab codes, supplementary videos as well as the analyzed three-dimensional tracking data of swarming mosquitoes is publicly available in the DRYAD repository: http://datadryad.org/stash/share/yyYPpUsKvRzBFxRI89pantIC9Fp19PP3YDpfPi9TZ6o.

## Supporting information titles and captions

**Table S1. List of all parameters used in our agent-based model.** Here, we provide a complete list of parameters and their default values as used in the agent-based model. The model description follows the ODD protocol (Grimm et al., 2020) in a simplified form. The model code is implemented in Matlab 2022b and in supplementary Code S1.

**Video S1. Example of the swarming flight kinematics of a single initiator mosquito.** Three-dimensional and two-dimensional views of the flight dynamic of the initiator male *Anopheles coluzzii* (i.e. the single first swarming mosquito), recorded in the lab above a black swarm marker of 40×40cm. This mosquito was flying in isolation.

**Video S2. Example of the flight kinematics of a single mosquito flying in a multi-individual swarm.** Three-dimensional coordinates, speed acceleration and angular speed of a single swarming *Anopheles coluzzii* male mosquito recorded in the lab above a black swarm marker of 40×40cm. This mosquito was flying in a multi-individual swarm.

**Video S3. Example of the swarming flight kinematics of multiple mosquitoes.** Three-dimensional and two-dimensional views of the flight kinematics of multiple male *Anopheles coluzzii* mosquitoes recorded in the lab above a black swarm marker of 40×40cm.

**Video S4. Example of the swarming flight kinematics of ten simulated mosquitoes.** Three-dimensional and two-dimensional views of the flight kinematics of ten simulated male mosquitoes above a black swarm marker of 40×40cm, using an agent-based model and minimally interactive behavioral rules (with collision avoidance).

**Database S1. Database of the experiment. Flight tracks of *Anopheles coluzzii* male mosquitoes swarming above a swarm marker in sunset simulated laboratory conditions.** A Matlab (all_data.mat) file containing three-dimensional tracks of all flying mosquitoes, described as the time t (s) and the spatial {x, y, z} (m) coordinates of the mosquito at each video frame. The coordinates are in meters, and in the world reference frame as defined in figure 1, with z oriented vertically up, and the origin of the coordinate frame at the center of the swarm marker. The trajectories were determined as described in the materials and methods.

**Database S2. Database of the simulation. Flight tracks of simulated male mosquitoes swarming above a swarm marker.** Two Matlab (all_data.mat) files containing three-dimensional tracks of all flying simulated mosquitoes with or without collision avoidance rule, described as the time t (s) and the spatial {x, y, z} (m) coordinates of the mosquito at each video frame. The coordinates are in meters, and in the world reference frame as defined in figure 1, with z oriented vertically up, and the origin of the coordinate frame at the center of the swarm marker. The trajectories were determined as described in the materials and methods.

**Code S1. Matlab analysis codes.** Contains all original Matlab codes that were written to perform the analysis of the article and to generate the figures.

