## Supplementary figures for "The complex swarming dynamics of malaria mosquitoes emerge from simple minimally-interactive behavioral rules"

**The PDF file includes:**

Figs S1 to S15

**Other Supplementary Materials for this manuscript include the following:**

Table S1
Videos S1 to S4
Database S1 and S2
Code S1

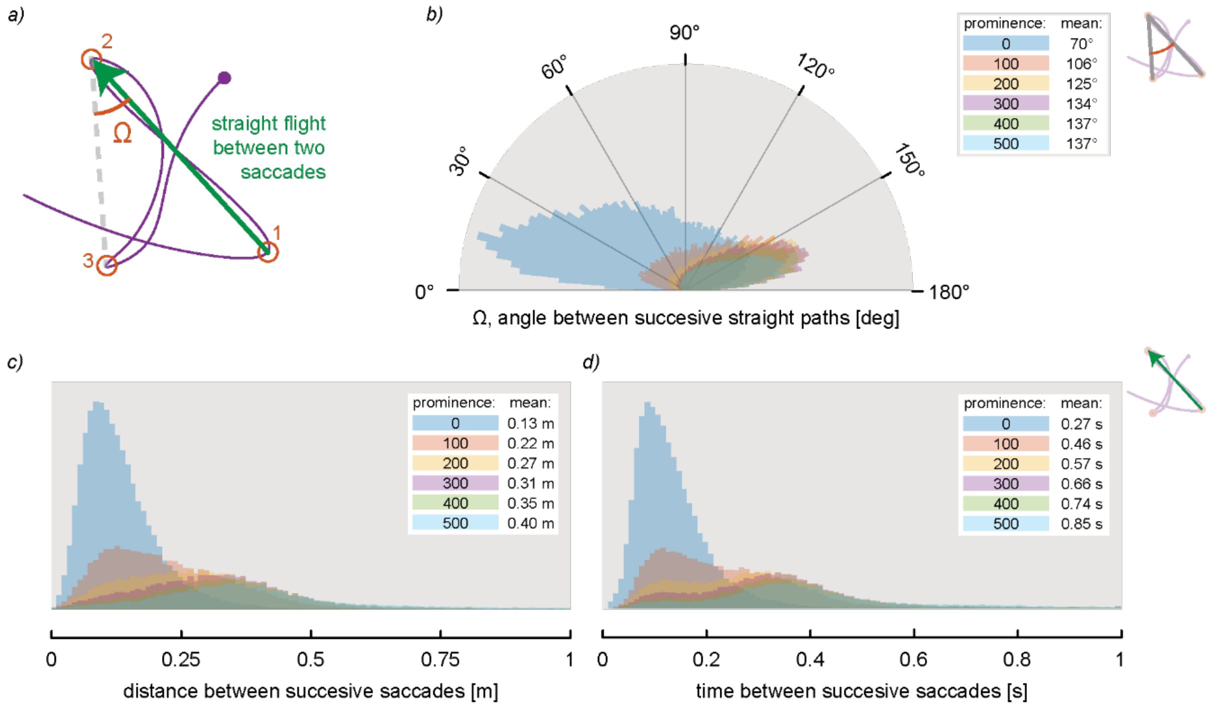


**Fig. S1. Sensitivity analysis to select prominence for saccadic turn detection.** Experimental swarming data are from the Poda dataset (Poda et al., 2024); see Fig S13 for equivalent data from the Feugère datasets. (*a*) Top view of a swarming flight trajectory, including three consecutive saccadic turns (1,2,3; defined in orange), the straight flight path between saccades (green) and the saccadic turn angle (W). (*b-d*) Distributions of the saccadic turn and straight flight metrics within all 938 recorded swarming flight tracks, for a range of prominence values (see legends, including mean values). (*b*) Polar histogram of the saccadic turn angle **Ω**. (*c-d*) Histograms of the distance travelled (*c*) and time duration (*d*) between successive saccades. A prominence of zero resulted in the detection of a large number of angular speed peaks, being associated with false positives. Based on these results, we selected a prominence threshold value of 300 for identifying saccadic maneuvers, and which is the same value as used for fruit flies (Van Breugel et al., 2022).

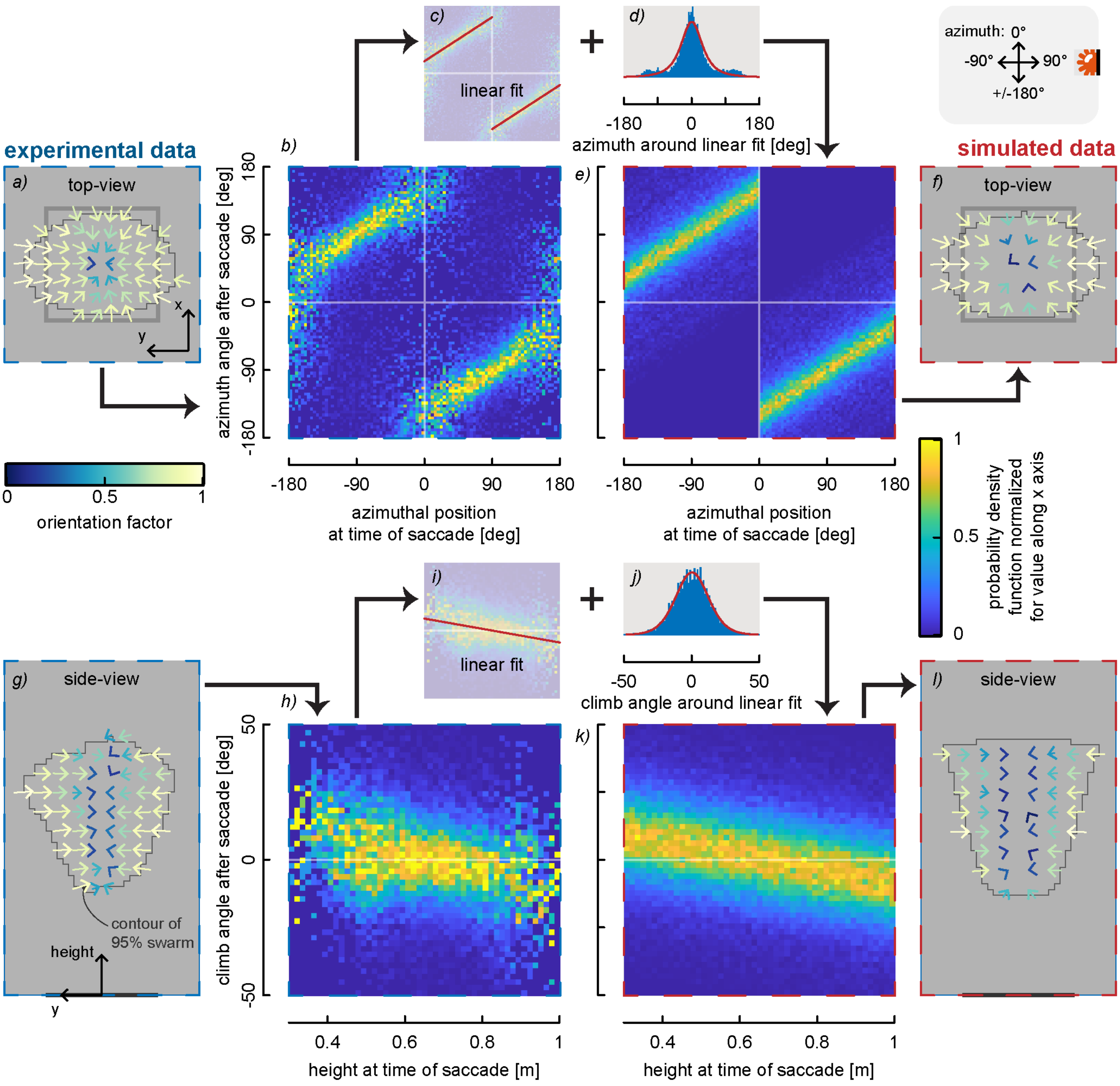


**Fig. S2. Flight direction after a saccadic turn, from both experimental Poda dataset and the simulation dataset.** All experimental data are shown on the left (*n*=938 tracks), and simulation data on the right (*n*=938 tracks), as indicated by the blue and red dashed boxes, respectively. (*a,g,f,l*) Top view (*a,f*) and side view (*g,l*) of the flight velocity vectors of mosquitoes after exhibiting a saccadic maneuver, from experiments (*a,g*) and simulations (*f,l*). The black bar and grey square represent the swarm marker. (*b,e,h,k*) Distributions of differences in azimuth angles (*b,e*) and climb angles (*h,k*) between two successive saccades, from experiments (*b,h*) and simulations (*e,k*). The simulated azimuth and climb angle distributions were constructed using linear fits (*c,i*) and corresponding distributions (*d,j*) of the experimental data.

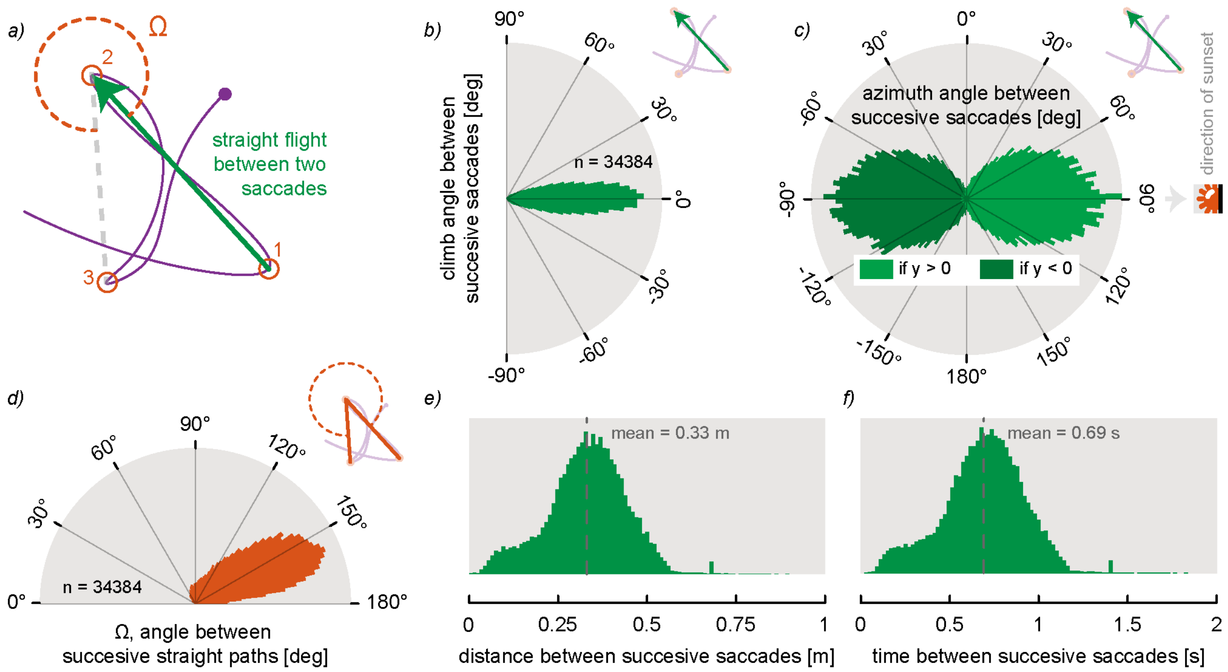


**Fig. S3. Distributions of turn kinematics of the in-swarm saccadic flight maneuvers.** Results are from the Poda dataset (Poda et al., 2024); see Fig S13 for equivalent results from the Feugère datasets (Feugère et al., 2022). (*a*) Top view of a swarming flight trajectory, including three consecutive saccadic turns (1,2,3; defined in orange), the straight flight path between saccades (green) and the saccadic turn angle (W). (*b,c*) Polar histograms showing the climb angle (*b*, angle out of the horizontal plane) and azimuth angle (*c*; angle within the horizontal plane) between successive saccades. Azimuth angles of -90° and +90° correspond directions away and towards the sunset, respectively. (*d*) Polar histogram of the saccadic turn angle **Ω**. (*e,f*) Histograms showing the distance traveled and time between successive peaks of angular speed for all 938 flight tracks.

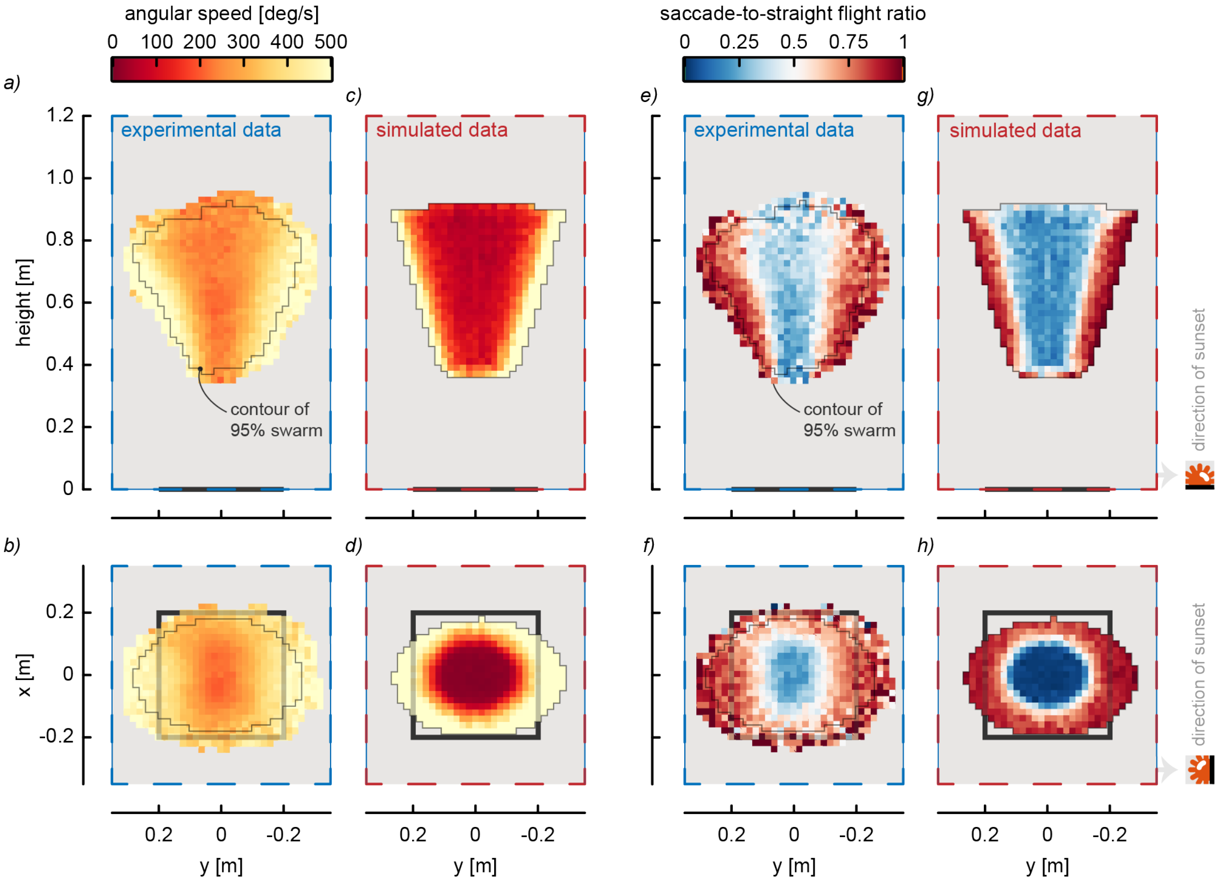


**Fig. S4. Comparison of the saccadic turn characteristics and distributions within the experimental Poda dataset and the agent-based simulation dataset.** See Fig S15 for equivalent data from the experimental Feugère datasets. The top and bottom rows show the side and top projection of the in-swarm distributions, respectively; experimental results (*n*=938 tracks) and simulated results (*n*=938 tracks) are highlighted by blue and red dashed boxes, respectively. (*a-d*) The angular speed distributions, within experimental swarms (*a,b*) and simulated swarms (c,*d*). (*e-h*) The distribution of the saccade-to-straight flight ratio, within experimental swarms (*e,f*) and simulated swarms (*g,h*). The thin grey line shows the contour of the volume comprising 95% of the three-dimensional tracking data, the black bar and square represent the swarm marker, and the sunset location is indicated on the right. Data was shown at a cell size resolution of 2x2cm. No data was shown for cells with less than an average of one track per recording period and per swarm (i.e. 3 recording periods * 6 swarms = 18).

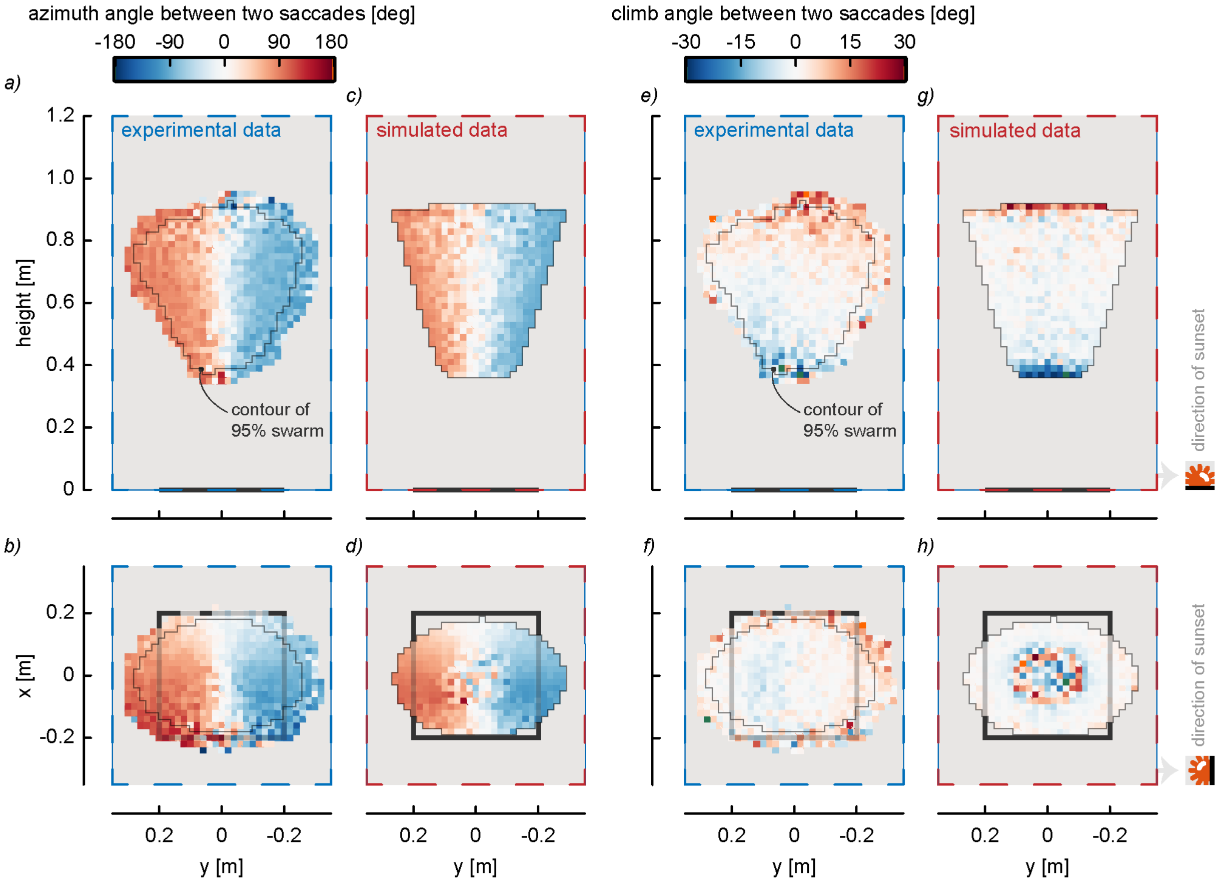


**Fig. S5. Comparison of the azimuth and climb angle distributions within the experimental Poda dataset and the agent-based simulation dataset.** The top and bottom rows show the side and top projection of the in-swarm distributions, respectively; experimental results ($n$=938 tracks) and simulated data ($n$=938 tracks) are highlighted by blue and red dashed boxes, respectively. (*a-d*) The azimuth angle distributions within experimental swarms (*a,b*) and simulated swarms (*c,d*). (*e-h*) The climb angle distributions within experimental swarms (*e,f*) and simulated swarms (*g,h*). All angles are angular differences between the straight flight paths before and after the saccadic turn. The thin grey line shows the contour of the volume comprising 95% of the three-dimensional tracking data, the black bar and square represent the swarm marker, and the sunset location is indicated on the right. Data was shown at a cell size resolution of 2x2cm. No data was shown for cells with less than an average of one track per recording period and per swarm (i.e. 3 recording periods * 6 swarms = 18).

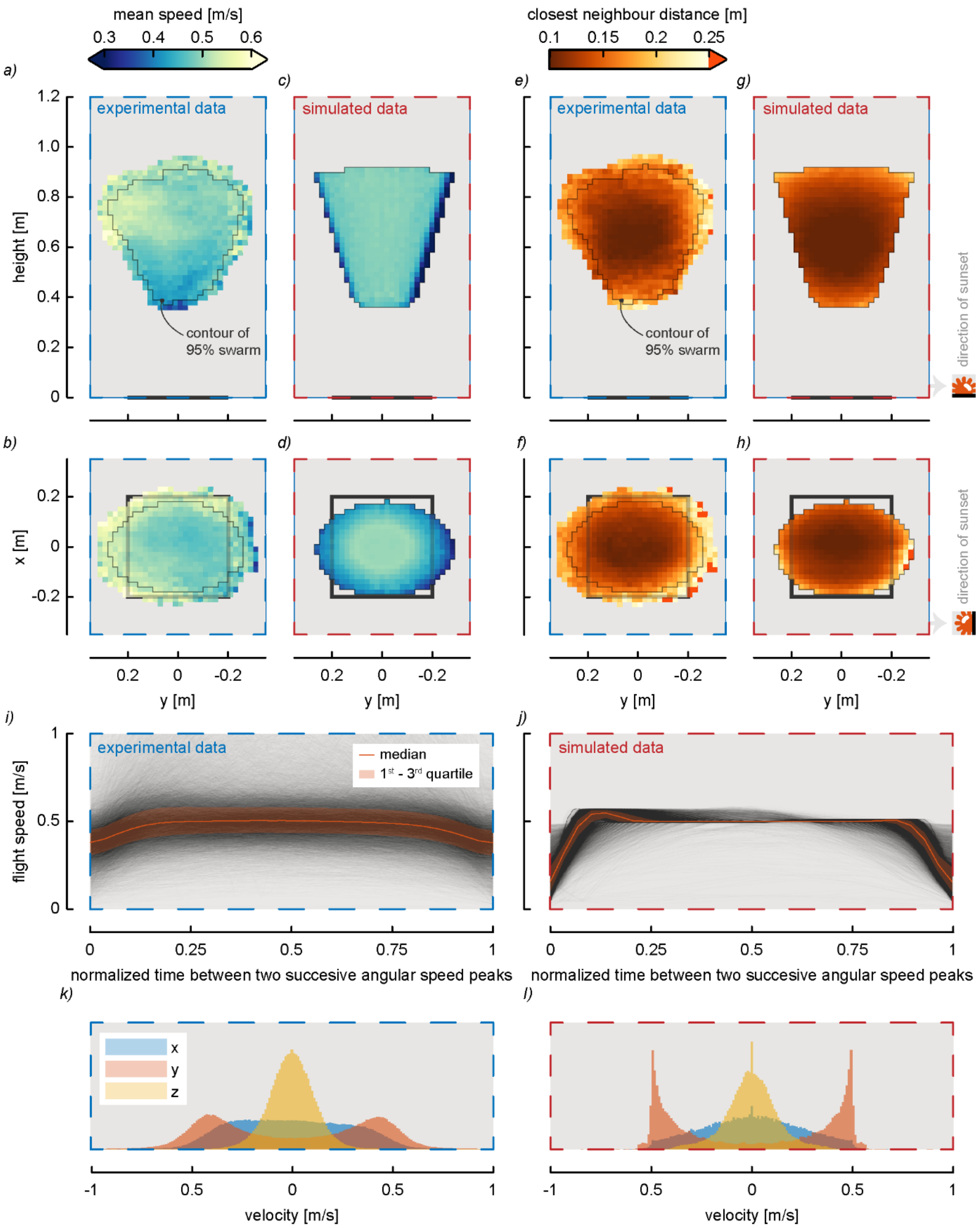


**Fig. S6. Comparison of the speed and density distributions within the experimental Poda dataset and the agent-based simulation dataset.** See Figs S12 and S14 for equivalent data from the experimental Feugère datasets. Experimental results (*n*=938 tracks) and simulated results (*n*=938 tracks) are highlighted by blue and red dashed boxes, respectively. (*a-d*) Side view (*a,c*) and top view (*b,d*) of the flight speed distributions within experimental swarms (*a,b*) and simulated swarms (*c,d*). (*e-h*) Side view (*e,g*) and top view (*f,h*) of the closest neighbor distance distribution within experimental swarms (*e,f*) and simulated swarms (*g,h*). (*a-h*) The thin grey line shows the contour of the volume comprising 95% of the three-dimensional tracking data, the black bar and square represent the swarm marker, and the sunset location is indicated on the right. Data was shown at a cell size resolution of 2x2cm. No data was shown for cells with less than an average of one track per recording period and per swarm (i.e. 3 recording periods * 6 swarms = 18). (*i,j*) Density plot of straight flight speed between two saccadic turns versus time normalized by the straight flight path duration, within experimental swarms (*i*) and simulated swarms (*j*). (*k-l*) histograms of the 3D flight velocity components within the experimental swarms (*k*) and simulated swarms (*l*).

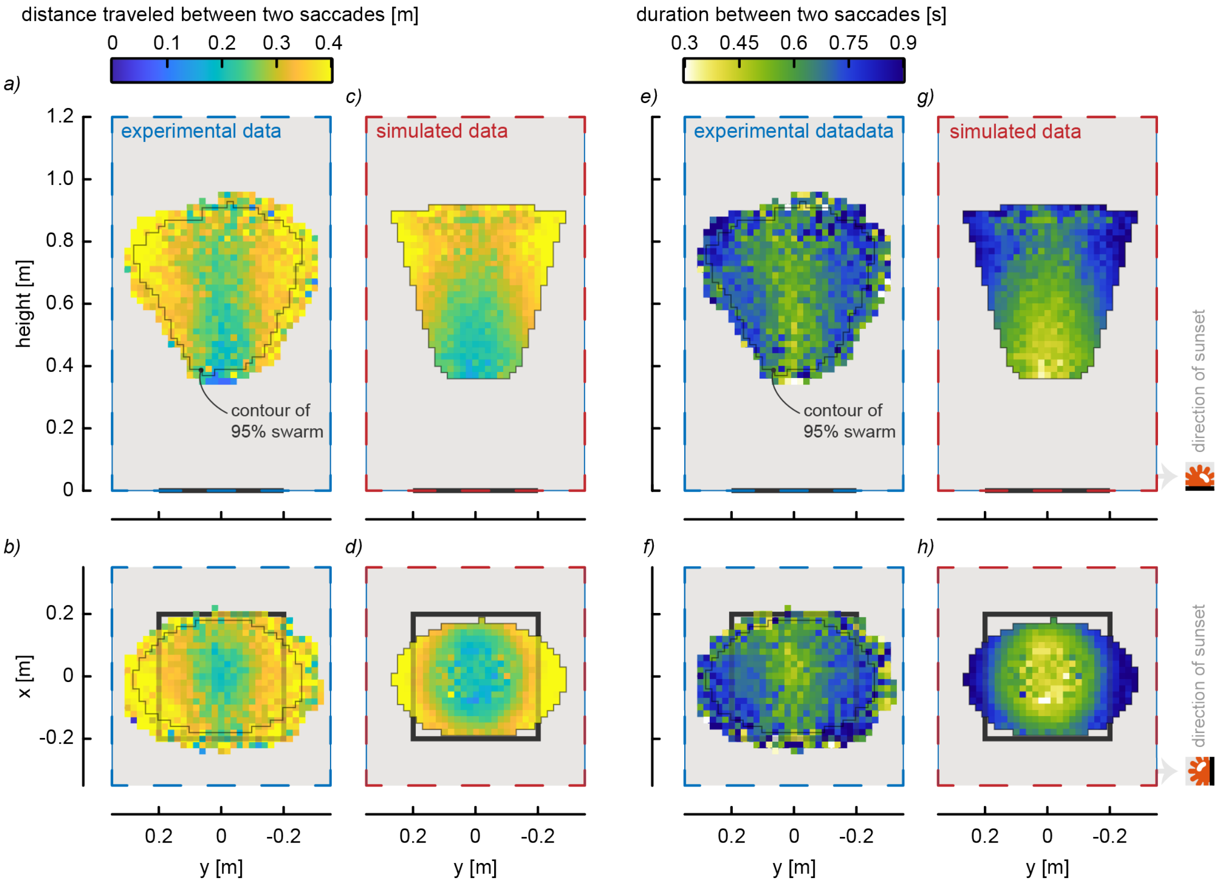


**Fig. S7. Comparison of the straight flight paths characteristics and distributions within the experimental Poda dataset and the agent-based simulation dataset.** The top and bottom rows show the side and top projection of the in-swarm distributions, respectively; experimental results ($n$=938 tracks) and simulated results ($n$=938 tracks) are highlighted by blue and red dashed boxes, respectively. (*a-d*) The distributions of straight flight distances, within experimental swarms (*a,b*) and simulated swarms (c,*d*). (*e-h*) The distribution of the straight flight durations, within experimental swarms (*e,f*) and simulated swarms (*g,h*). The thin grey line shows the contour of the volume comprising 95% of the three-dimensional tracking data, the black bar and square represent the swarm marker, and the sunset location is indicated on the right. Data was shown at a cell size resolution of 2x2cm. No data was shown for cells with less than an average of one track per recording period and per swarm (i.e. 3 recording periods * 6 swarms = 18).

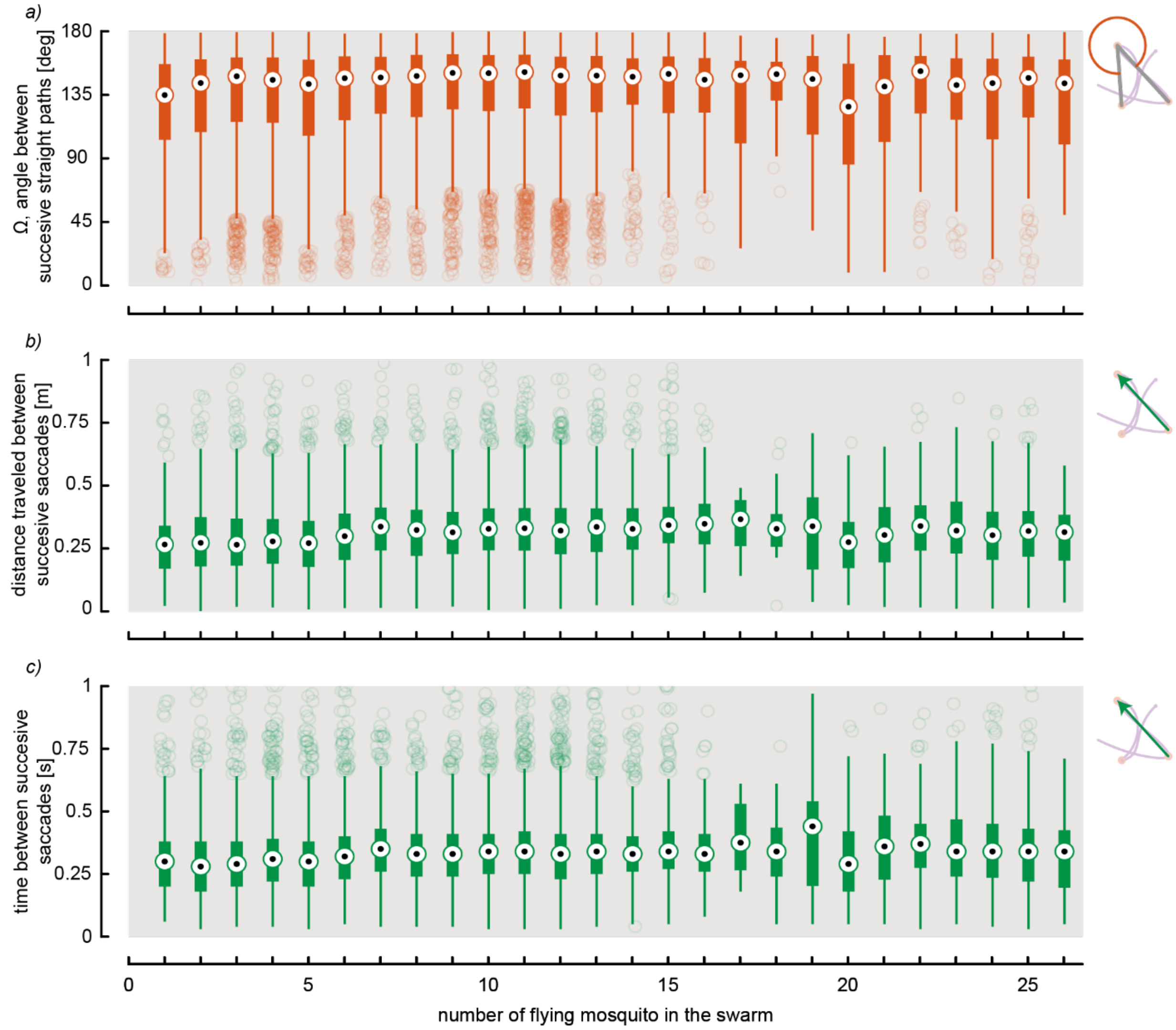


**Fig. S8. We observed no effect of the number of swarming mosquitoes on the individual flight kinematics.** Results are from the Poda dataset. The primary swarming metrics as a function of the number of mosquitoes in the swarm, throughout all three recordings within six swarms. During these recordings, the number of swarming individuals ranged from a single swarm initiator to 26 individuals. The primary swarming metrics include the saccadic turn angle Ω (*a*), the distance travelled during the straight flight phase between successive saccades (*b*), and the time duration of these straight flight phases (*c*). The saccadic turn angle and straight flight phase are defined in the flight track on the right, as the orange angle and green vector, respectively.

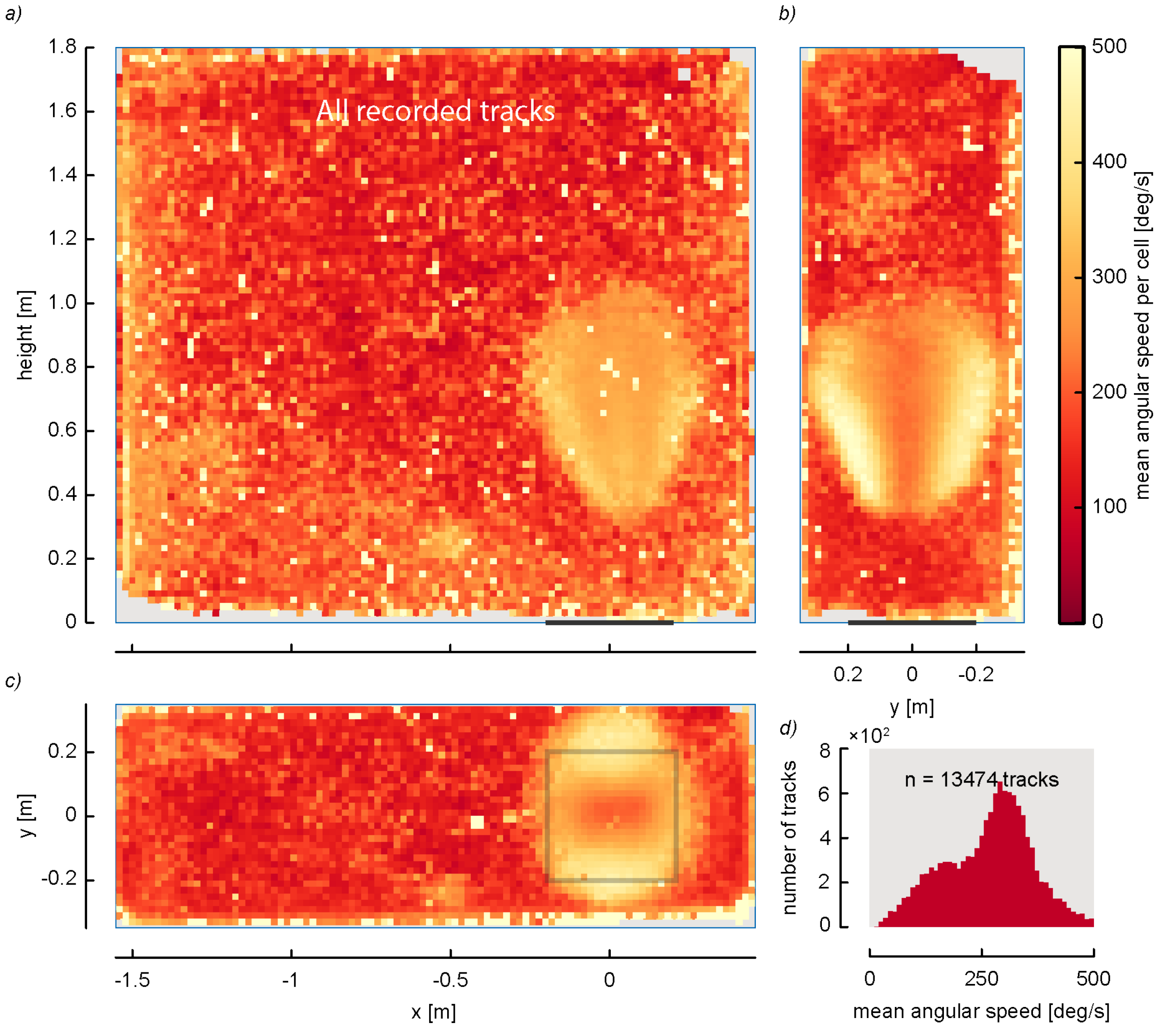


**Fig. S9. Spatial distribution of angular speed for all recorded flight tracks in the Poda dataset, including tracks of non-swarming mosquitoes.** Front view (*a*), side view (*b*) and top view (*c*) of the spatial distribution of the angular flight speeds (*n*=938 tracks). Data was shown at a cell size resolution of 2x2 cm. The swarm marker is represented by black line (*a*,*b*) or black square (*c*). The angular speeds are higher within the swarm volume than outside. (*d*) Histogram of the mean angular flight speeds.

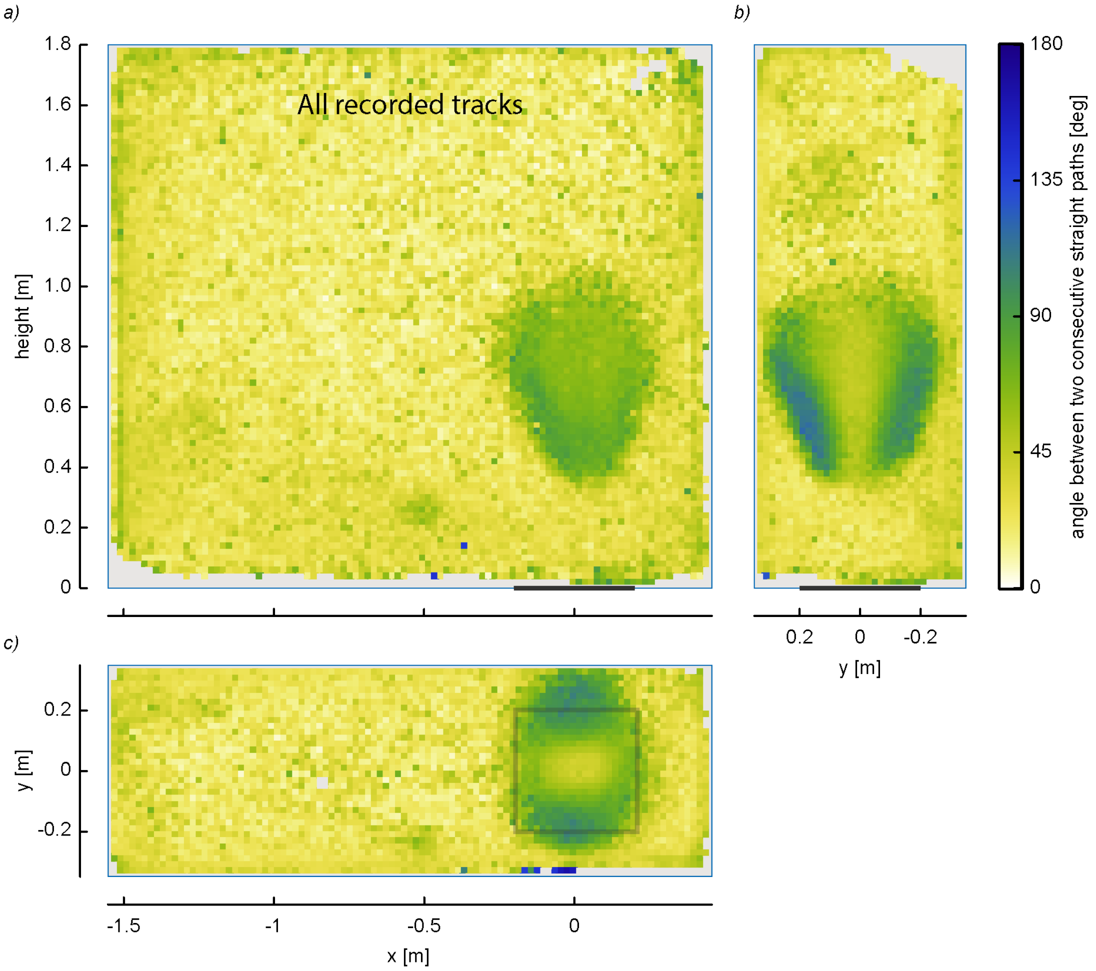


**Fig. S10. Spatial distribution of the angle between two successive saccades for all recorded flight tracks in the Poda dataset, including tracks of non-swarming mosquitoes.** Front view (*a*), side view (*b*) and top view (*c*) of the spatial distribution of the saccadic turn angle Ω ($n$ = 19,572 straight flight phases within 938 flight tracks). Data was shown at a cell size resolution of 2x2cm. The swarm marker is represented by black line (*a*,*b*) or black square (*c*). The saccadic turn angles are higher within the swarm volume than outside.

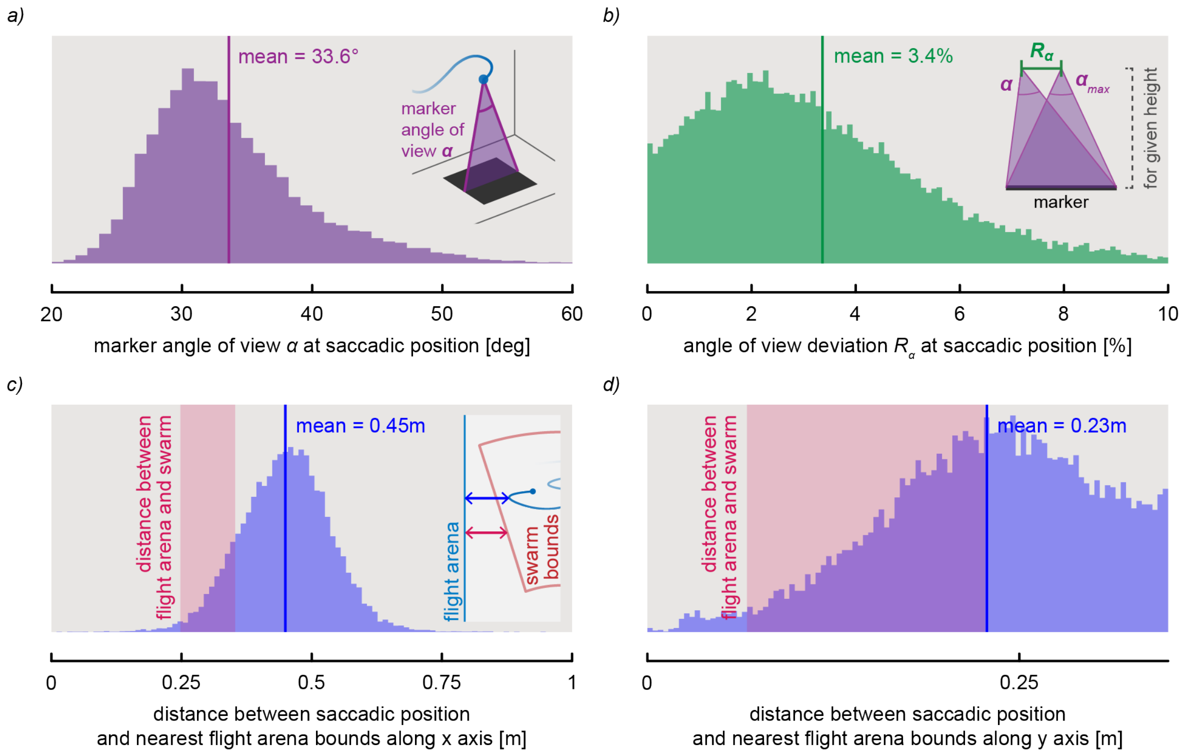


**Fig. S11. Environmental cues experienced during the in-swarm saccadic turns.** Results are from the Poda dataset. (*a-b*) Histograms of the viewing angle of the swarm marker *α* (*a*) and viewing-angle deviation $R_α$ (*b*), at each saccadic turn position (*n* = **1**9,572 saccades within 938 flight tracks). The mean viewing-angle deviation ($R_α$=3.4%) is below the 4.5% threshold defining the swarm boundary, indicating that mosquitoes typically turn back toward the swarm center before reaching the swarm limit. (*c-d*) Histograms of the distance from the arena wall at each saccadic turn, along both the x-axis (*c*) and y-axis (*d*). The distribution of distances to the arena walls at swarm boundary are shown in red, where shortest and longest distance are at the top and bottom of the swarm (see schematic in (*c*)). The mean wall distance for all saccades combined were respectively 45 cm and 23 cm along the *x*-axis and *y*-axis, which equals ~75 and ~38 body lengths. By contrast, documented wall-interaction distances in mosquitoes are on the order of only one to a few body lengths (Cribellier et al., 2024; Nakata et al., 2020).

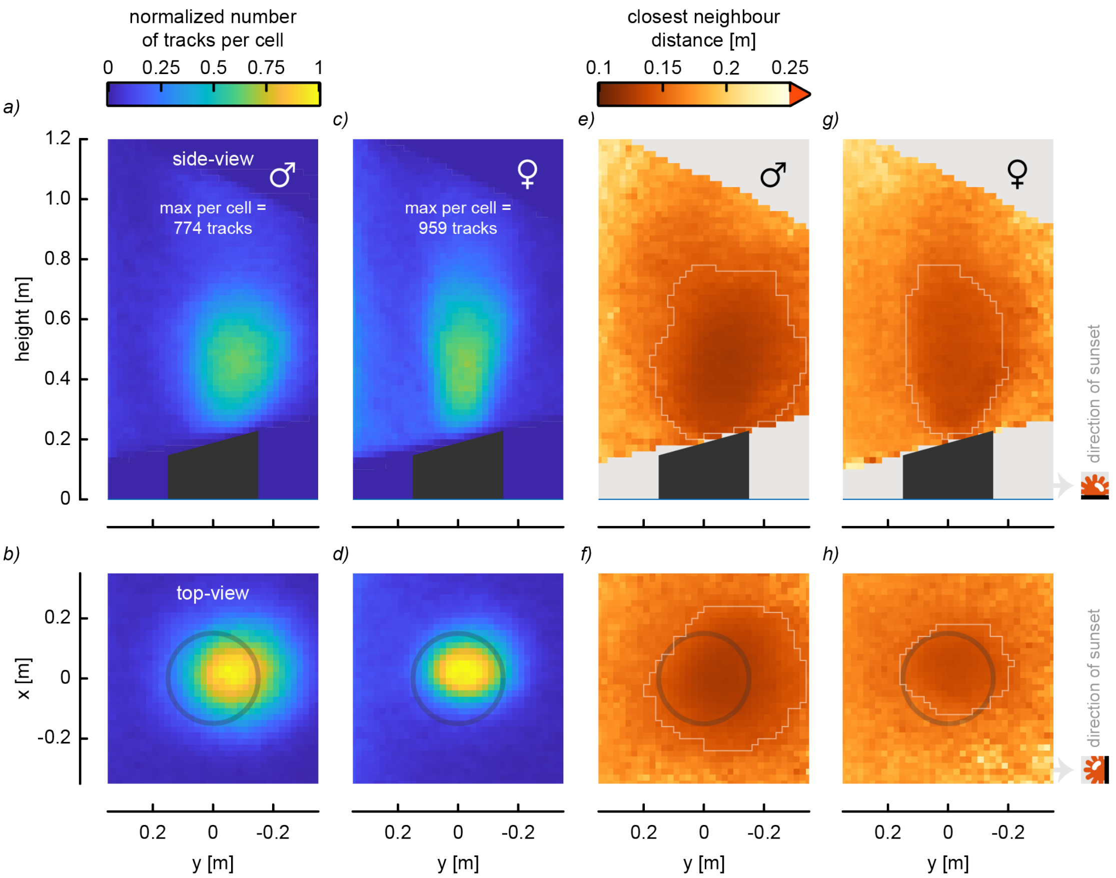


**Fig. S12. Positions, shapes and densities of swarms produced by male and female *An. coluzzii* mosquitoes from the Feugère dataset (Feugère et al., 2022).** The top and bottom rows show the side and top projection of the distributions, respectively. (*a-d*) Normalized track densities for swarming males (*a,b*; $n$ = 3121 trajectories) and females (*c,d*; $n$ = 5933 trajectories). (*e-h*) Mean distance between closest neighbors (*e-h*), an indirect measurement of swarm density, for swarming males (*e,f*) and females (*g,h*). The white contour line defines the 3D swarming regions, where the mean distance between closest neighbors was lower than the marker diameter (0.3 m).

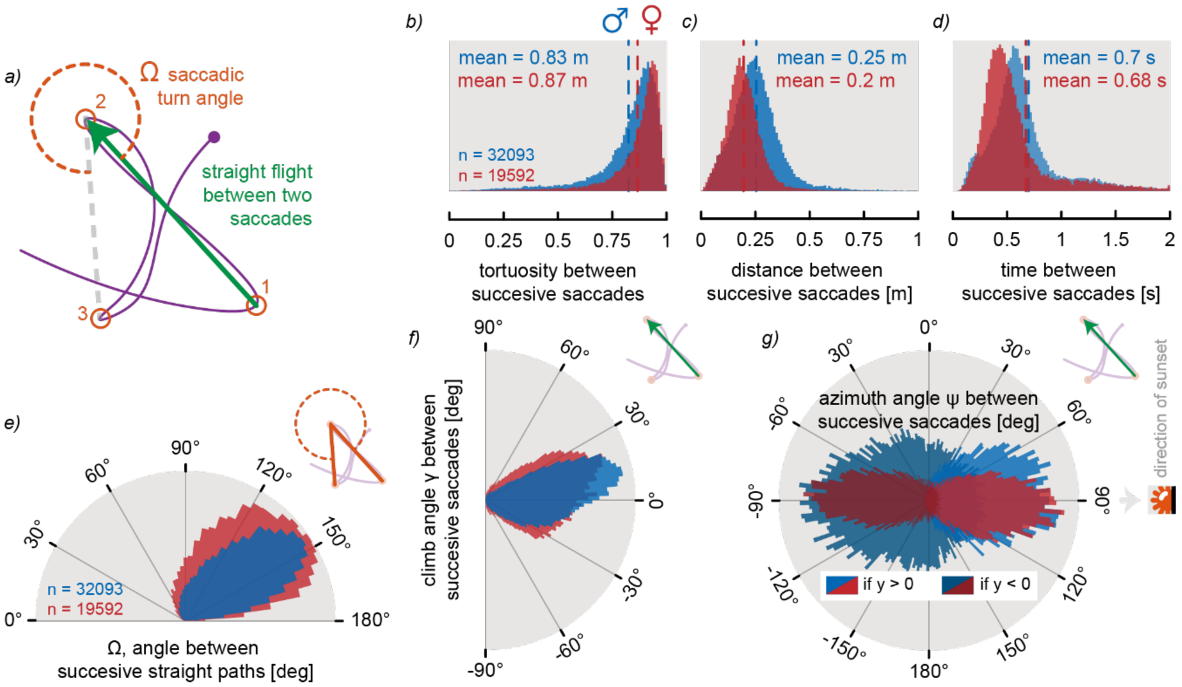


**Fig. S13. Distributions of turn kinematics of the in-swarm saccadic flight maneuvers, for swarming male and female *An. coluzzii* from the Feugère dataset (Feugère et al., 2022).** See Figs 2, S1 and S3 for equivalent results from the Poda dataset. (*a*) Top view of a swarming flight trajectory, including three consecutive saccadic turns (1,2,3; defined in orange), the straight flight path between saccades (green vector) and the saccadic turn angle (Ω). (*b-d*) Histograms showing the tortuosity (*b*), distance traveled (*c*), and time duration (*d*) of the straight flight phases between successive saccadic turns of males and females mosquitoes (*n*=32,093 and 19,592 straight flight phases within 1848 and 1740 trajectories, respectively). (*e*) Polar histogram of the saccadic turn angle Ω. (*f,g*) Polar histograms of the climb angle (i.e. angle from the horizontal plane; *f*) and azimuth angle (i.e. angle on horizontal plane; *g*) during the straight flight phases. Azimuth angles of -90° and +90° correspond to directions away and towards the sunset, respectively (see sunset symbol in *g*). Flight dynamics of both males and females mosquitoes are very similar to those of the Poda dataset (Fig. 2). Notable minor differences are: 1) climb angles of males and females were less concentrated around 0° than in our own experiments, maybe because mosquitoes flew on a similar tilted plan than the sloped top edge of cylindrical marker, but both dataset show a strong preference for horizontal movements; 2) males show higher diversity of azimuth angles than both the females and the males in the Poda dataset.

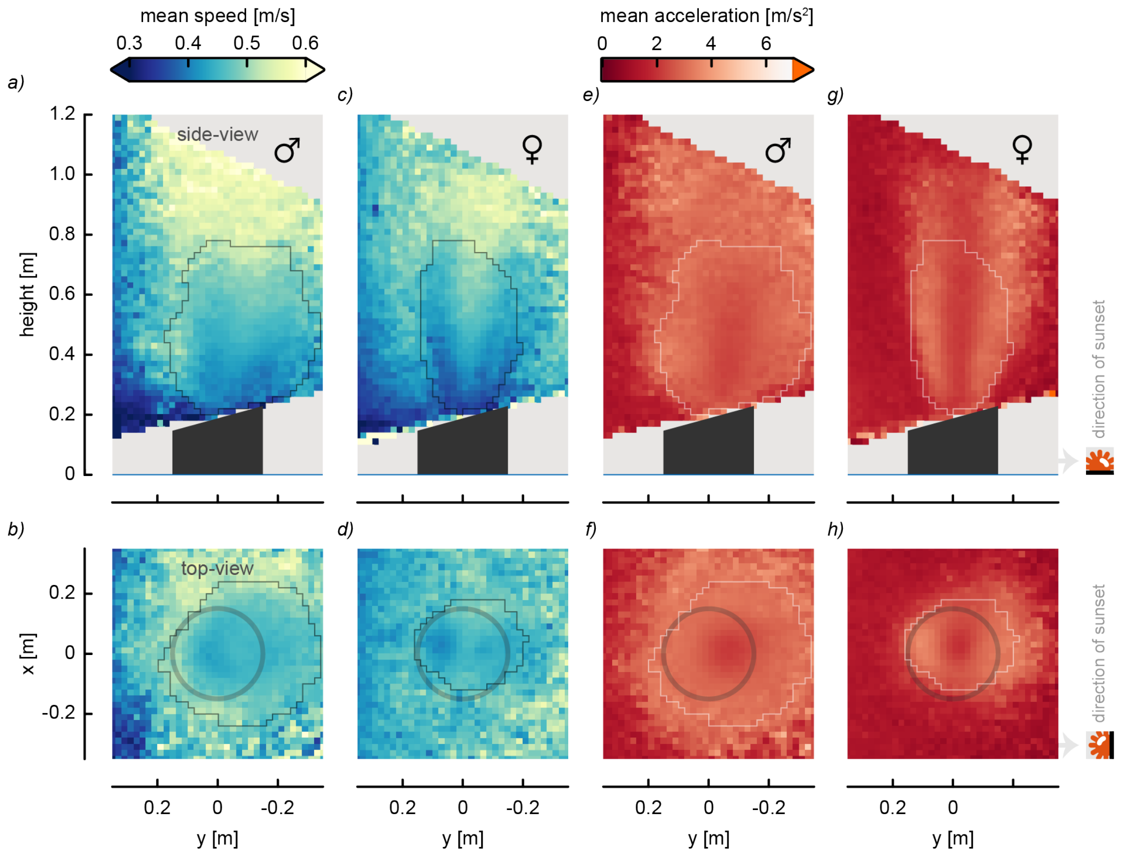


**Fig. S14. Spatial distribution of flight speeds and accelerations of swarming *An. coluzzii* males and females from the Feugère dataset (Feugère et al., 2022).** The top and bottom rows show the side and top projection of the distributions, respectively. (*a-d*) Normalized track densities for swarming males (*a,b*) and females (*c,d*). (*e-h*) Mean distance between closest neighbors (*e-h*), an indirect measurement of swarm density, for swarming males (*e,f*) and females (*g,h*). See Fig S6 for equivalent results from the Poda dataset. (Feugère et al., 2022)

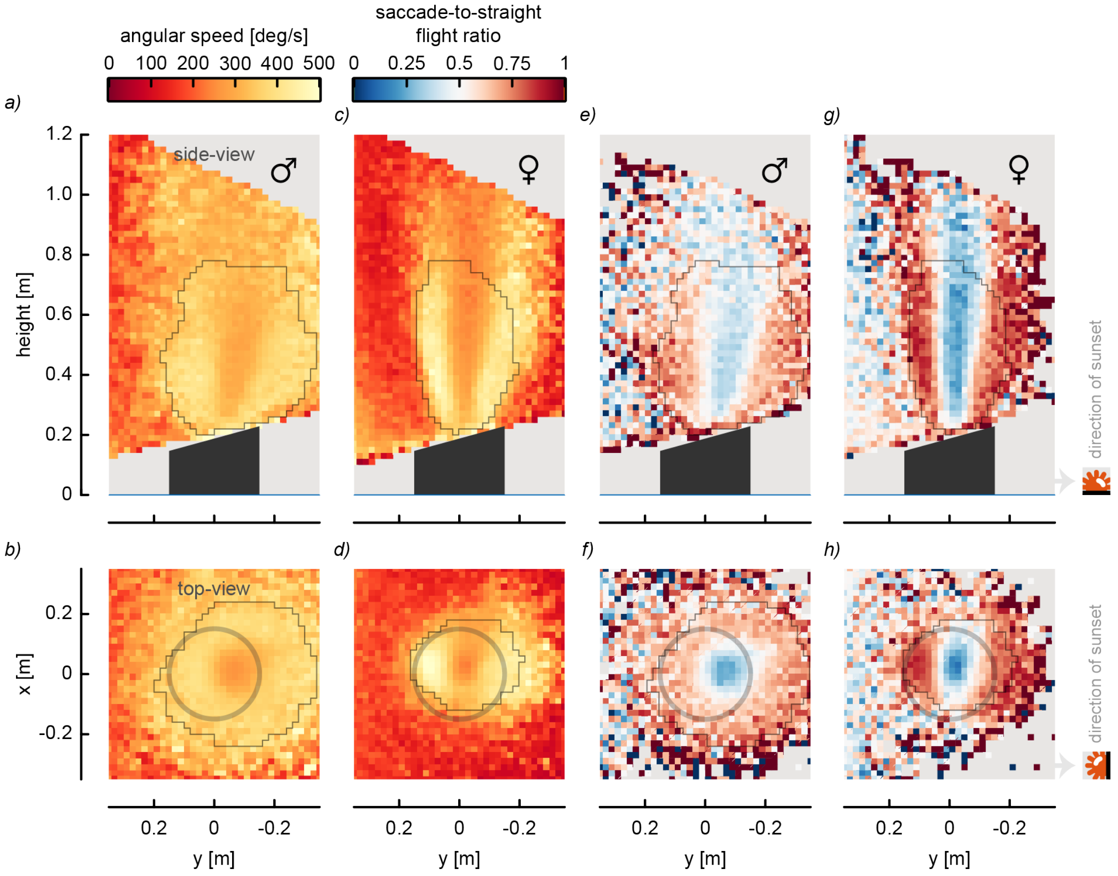


**Fig. S15. Saccadic maneuvers spatial distribution of swarming *An. coluzzii* males and females from the Feugère dataset (Feugère et al., 2022).** The top and bottom rows show the side and top projection of the distributions, respectively. (*a-d*) The normalized track densities for swarming males (*a,b*) and females (*c,d*). (*e-h*) Mean distance between closest neighbors, an indirect measurement of swarm density, for swarming males (*e,f*) and females (*g,h*). These flight dynamics are similar to those for the Poda dataset (Figs 3 and S4), namely saccadic maneuvers at the edge of the swarm and straight flight path through the middle of the swarm, with a flight direction preference along sunset axis (*f-h*). This is particularly apparent for female swarms (*g,h*).
